# Synthetic Data to Explore Transcriptional Regulation of Differentially Expressed Genes in Ovarian Cancer

**DOI:** 10.64898/2026.01.15.699618

**Authors:** Zhang Shuo, Kumar Selvarajoo

## Abstract

In major diseases like cancer, many genes are dysregulated. While differential expression (DE) analyses help identify these genes, they do not reveal the transcriptional regulatory landscape between control and disease states. Here, we developed a process-driven model based on generalized stochastic transcriptional regulation (TR) to study DE genes in ovarian cancer. We generated over 39,000 synthetic gene expression profiles based on RNA-seq dataset of 11 samples (5 control FTE and 6 ovarian cancer), and fitted the model parameters to real data achieving between 82% and 99% transcriptome-wide similarity through various data analytics. Through the model we investigated the top 100 DE genes’ transcriptional regulation. Notably, *GSTA3*, *SNTN*, *DNAI2* and *KIF19*, which have been associated with cancer, possess significantly larger transcriptional quantal and RNA degradation rates, leading to their greater variability in cancer. This mechanistic understanding could potentially guide gene-targeted interventions for cancer or other diseases in the future.

## Introduction

Since the turn of the millennium, biological communities have been overwhelmed by a flood of high-throughput datasets across single and multi-omics scales. This has given rise to the fields of bioinformatics, computational systems biology, and biomedical data science. More recently, artificial intelligence in biology and health research has gained significant momentum [1,2]. These modern, interdisciplinary approaches are essential for interpreting large-scale datasets and for unravelling the complex, multi-dimensional, and multi-modal nature of biological and disease processes [3].

Transcriptome- and genome-wide expression data generation and analysis have made remarkable advances across all areas of biological research. From the early days of microarray-based gene expression profiling, the development of sophisticated next-generation sequencing techniques has provided invaluable insights, enabling a more comprehensive understanding of differences between control and test samples. Notably, differential gene expression analyses have revealed not only the most important genes that vary between control (e.g., normal) and test (e.g., disease) samples, but also their associated biological networks and functions [4,5,6]. This approach allows for the study of disease processes as interconnected and co-regulated network structures [7]. More recently, single-cell analyses have revealed gene expression variations within complex multi-cellular environments, such as the tumor microenvironment, even within the same tissue samples [8,9,10]. Coupled with advancements in imaging technologies, modern spatial transcriptomics now offers detailed insights into heterogeneous gene expression patterns across cellular space and time [11].

Despite significant progress in characterizing correlated gene networks and uncovering gene expression variabilities, the mechanistic understanding of genome- or transcriptome-wide expression regulation remains largely underexplored. In addition to technical noise, biological noise arising from intrinsic and extrinsic factors further complicates the interpretation of gene expression and its variability [12, 13]. Mechanistic models, particularly those based on a dynamical systems approach, are crucial for interpreting differential behaviors and estimating noise [14,15]. For instance, such models have been employed to rationally design single-molecule quantification experiments for mouse glutamatergic neurons [16], as well as to uncover transcriptional amplification and quantal behaviors that contribute to significant variability between the 2- to 8-cell stages in both human and mouse developmental cells [17]. Therefore, a systems biology approach can provide valuable insights into the complex, transcriptome-wide regulatory landscape.

In this study, we applied a transcriptional modeling approach to investigate the RNA-seq expression profiles of control (fallopian tube epithelium cells, FTEs) and test (high-grade serous ovarian cancer cells, HGSOCs) [18]. We began by performing statistical and data analyses, including Pearson Correlation Coefficient [19,20,21], Shannon entropy [22,23], PCA, and noise, or squared Coefficient of Variation, [17,24]. Next, we developed stochastic transcriptional models, based on the Gillespie algorithm (incorporating intrinsic Poisson noise), to synthetically simulate the entire transcriptomes of FTEs and HGSOCs. Additionally, statistical (Gaussian) noise was integrated to account for the extrinsic noise observed in real cells [12]. Using these models, we then determined the model parameters by fitting the gene expression data to the real data for all genes. We conducted further data analyses to assess the overall quality of the synthetic transcriptomes generated by the models in comparison to the real data. The models provide mechanistic insights into how each gene is regulated between FTEs and HGSOCs, including predictions of the regulatory mechanisms governing differentially expressed (DE) genes (Figure 1). Overall, this work is intended to highlight the importance of using mechanistic models for generating transcriptome-wide synthetic data. Such models can enhance our understanding of each gene or a group of genes’ transcriptional regulation, particularly for future RNA-based therapeutic interventions.

**Figure 1.**
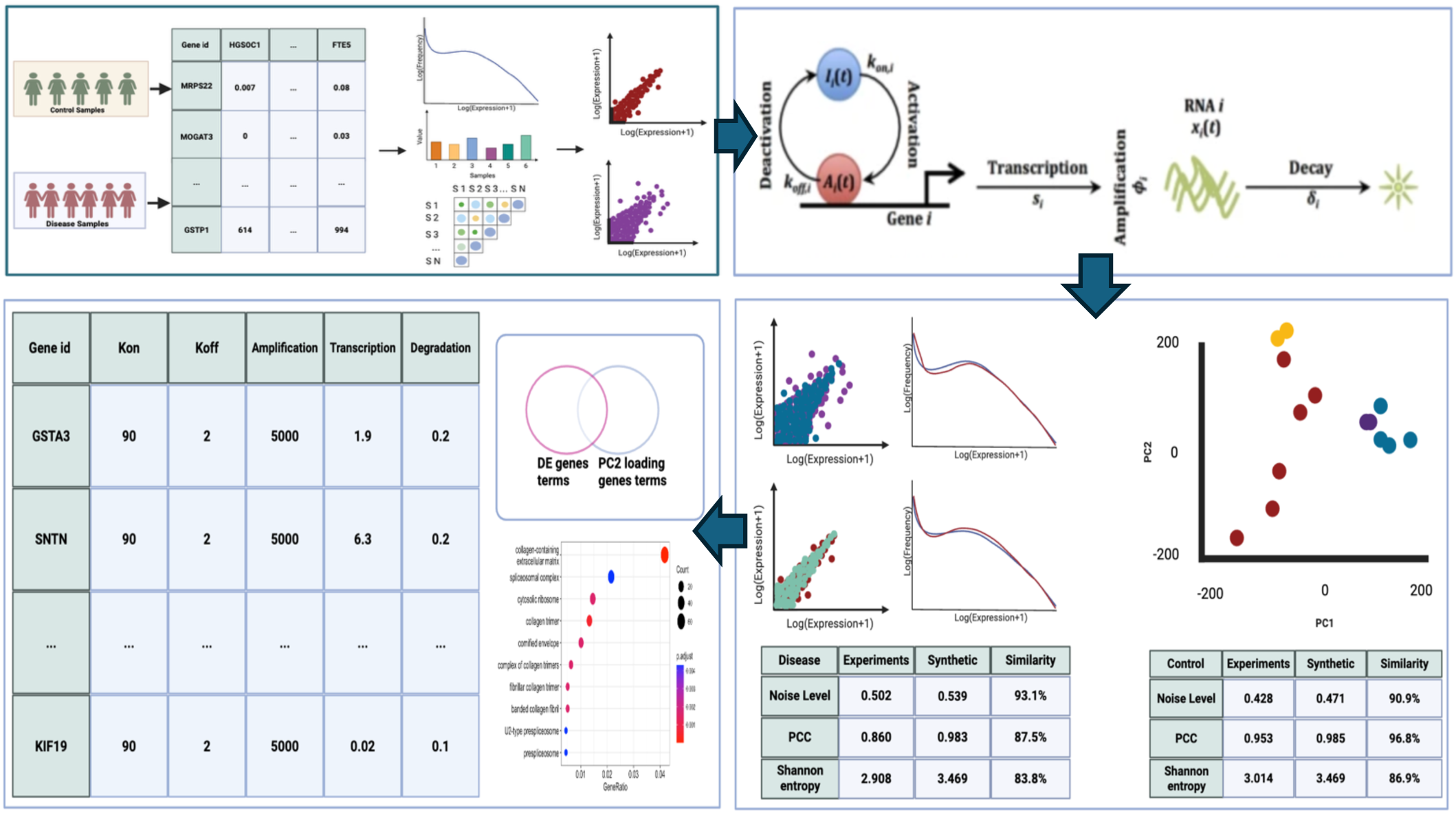
Schematic of Synthetic Data Generation Workflow. Step 1: The RNA-Seq (ovarian cancer) dataset is analysed using various data analytics. Step 2: A stochastic transcriptional model is developed to simulate and fit actual transcriptome-wide gene expressions data for all samples. The model parameters are varied for each sample to generate the synthetic data for its respective sample. Step 3: Simulated synthetic data are compared with real datasets through data analytic similarity tests. Step 4: Once synthetic data quality is assured, the TR mechanisms of DE genes are revealed through the interpretation from the model parameters.

## Result

### Ovarian Cancer Dataset

We focused on cancer-related datasets that included both control and normal counterpart cells for comparison. We selected datasets from the GEO database (GSE190688), which consist of RNA-seq data from 6 high-grade serous ovarian cancer (HGSOC) and 5 fallopian tube epithelium (FTE) samples collected from patients and donors [18]. Of the 48,162 genes available in the datasets, we excluded genes with zero expression across all samples, leaving 39,761 genes for analysis.

### Statistical Distributions, Scatter plots and Correlations

We plotted the gene expression distributions for all samples, which revealed bimodal patterns with a large number of genes showing either zero or low expression (Figure 2A for mean distribution, and Supplementary Fig. S1 upper table for all samples). Although the mean distributions of FTEs and HGSOCs illustrated a relatively similar pattern in the low to middle expressed region, HGSOC displayed a larger number of high-expressed genes (Figure 2A insert, and Fig. S1 upper table). To explore how gene expression patterns differ among replicates, we performed statistical fitting of the data which showed the estimated log-normal distributions for all replicates. (Fig.S1 lower table and Table 1).

**Figure 2.**
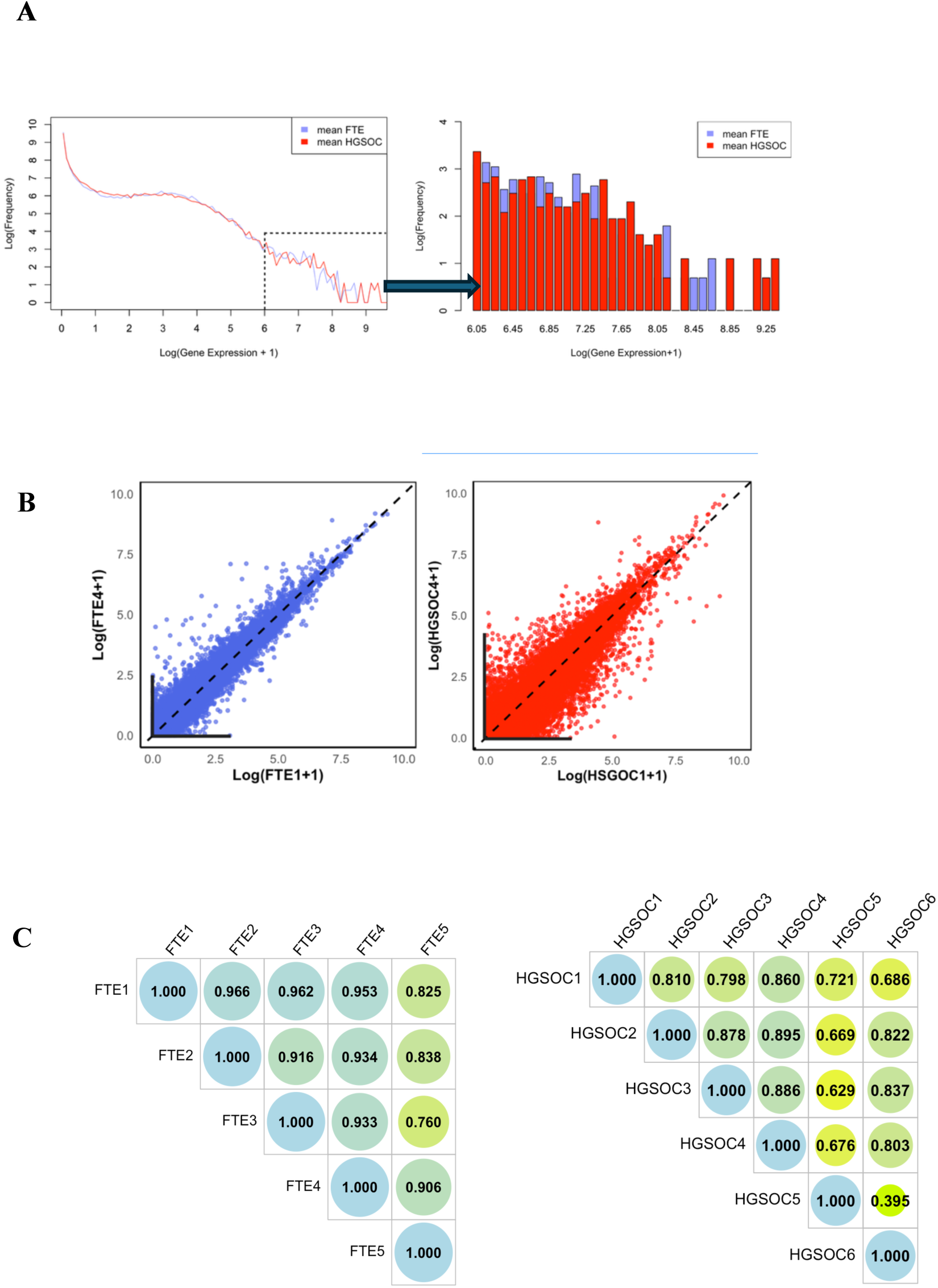
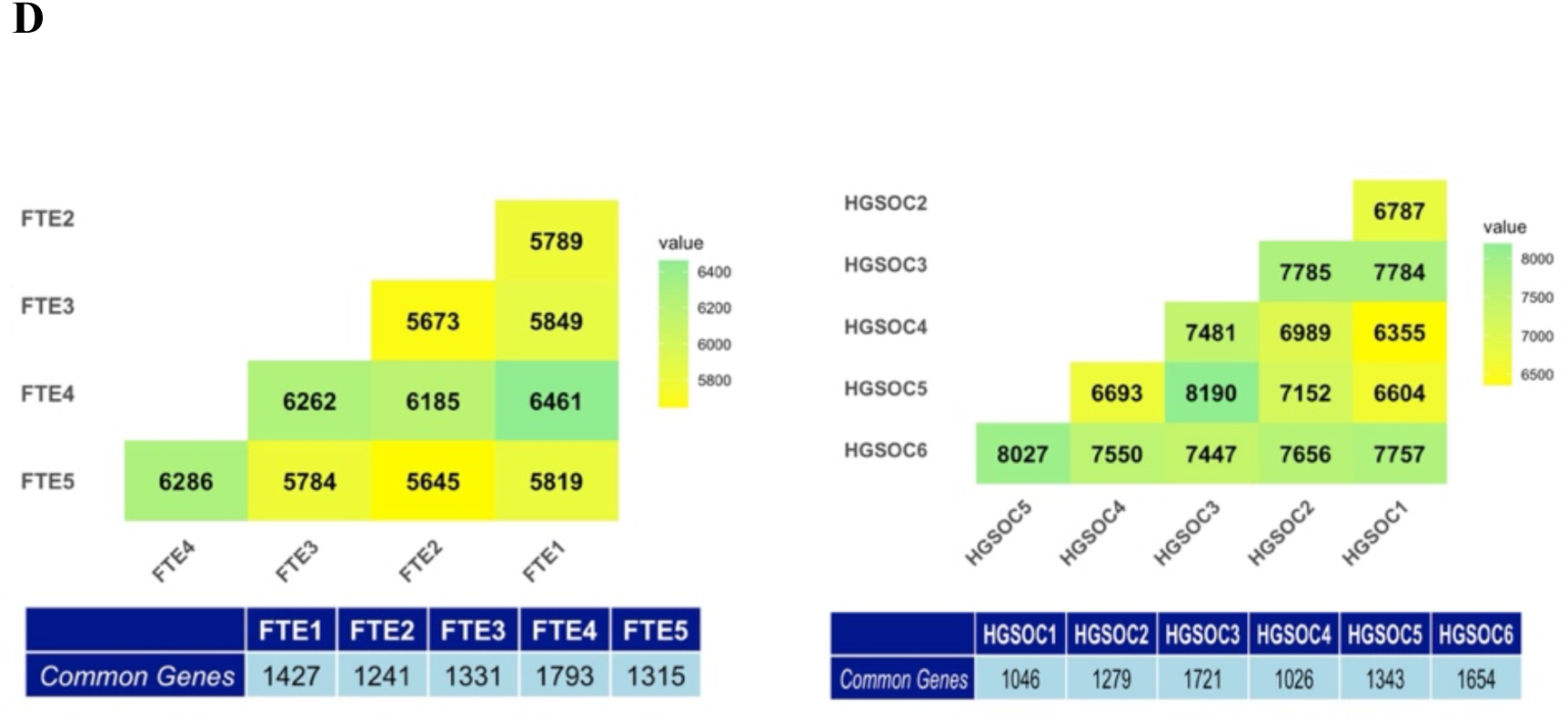
Data analytics to show statistical properties of FTEs and HGSOCs. (A) The mean gene expression distributions of FTEs (blue) and HGSOCs (red), see Supplementary Fig. S1 for all samples. Log-scale is applied to the gene expressions and frequency of distributions. Insert: the distribution plot of highly expressed regions of FTEs (blue) and HGSOCs (red), where the purple bars indicate the overlap of FTEs and HGSOCs. (B) The transcriptome-wide scatter plots of FTEs (blue) and HGSOCs (red), see Supplementary Fig. S2 for all samples. Log-scale is applied to the scatter plots and the toggle genes from the L-shape of the scatter plots are represented in black lines. (C) The bubble plots of Pearson correlation values for FTEs (left) and HGSOCs (right). (D) The distance matrix of the number of toggle genes for FTEs (top left) and HGSOCs (top right), and the number of toggle genes of one sample shared with other samples’ genes for FTEs (bottom left) and HGSOCs (bottom right).

**Table 1.** Variable parameter values for FTE and HGSOC samples.

| FTE replicate | $\mu$ (log-normal) | $\sigma$ (log-normal) |
| --- | --- | --- |
| FTE1 | 0.638 | 2.498 |
| FTE2 | 0.611 | 2.523 |
| FTE3 | 0.601 | 2.533 |
| FTE4 | 0.588 | 2.529 |
| FTE5 | 0.674 | 2.496 |
| HGSOC1 | 0.618 | 2.503 |
| HGSOC2 | 0.630 | 2.374 |
| HGSOC3 | 0.631 | 2.450 |
| HGSOC4 | 0.691 | 2.458 |
| HGSOC5 | 0.658 | 2.531 |
| HGSOC6 | 0.741 | 2.258 |

The identification of numerous zero-expressed genes in our datasets indicates the presence of toggle genes, a phenomenon previously observed in various cell types [25,26]. To investigate further, we created scatter plots comparing samples within the same cell condition (Figure 2B and Fig. S2). Notably, the transcriptome-wide scatter plots for FTE samples displayed a generally linear relationship, with expression values largely aligning along the identity line (Figure 2B, left, and Fig. S2). In contrast, the scatter plot for HGSOC samples showed greater dispersion from the identity line, with a notably higher number of variable genes between the same samples (Figure 2B, right, and Fig. S2).

Next, we computed the Pearson correlations for all samples and found that the correlations for FTE samples (∼0.90) were significantly higher than those for HGSOC samples (∼0.75) (Figure 2C) [19,20,21]. These results are expected, as cancer cells are typically more variable and heterogeneous [27]. Notably, the presence of toggle genes is evident in all datasets (Figure 2B, L-shaped black lines), with the total number of toggle genes between individual replicates and the common toggle genes across all samples presented (Figure 2D). Overall, HGSOCs had more toggle genes than FTEs, though fewer were shared across samples (Figure 2D, bottom). Thus, HGSOCs exhibited a greater number and variability of toggle genes.

### PCA, Shannon Entropy and Pair-wise Noise

Next, we used PCA as a dimensionality reduction technique to compare the different samples, which showed the first two components collectively accounted for about 41% of the dataset variance. It is obvious from the plot that FTE samples are clustered together, while the HGSOC samples are more spread out across the plot. PC1 separates the FTE samples (positive values) from the HGSOC samples (negative values), while PC2 captures the variability between the samples. In other words, the high variability among the HGSOCs is reflected by PC2.

To quantify the gene expression variability between samples, we computed Shannon entropy, a measure often used to assess the “disorder” within a sample’s gene expressions for all samples [17,22,23]. Although the analysis revealed similar mean Shannon entropy values (∼3.55 for FTE and ∼3.47 for HGSOC), the standard deviation, however, was noticeably higher for HGSOCs (Figure 3B, left), indicating more diversity in the gene expressions within the latter. Next, we used the square of the coefficient of variation, or noise, approach to compare sample-wise variability [17, 24]. The transcriptome-wide average noise for all pair-wise combinations of HGSOCs was significantly higher (0.62) compared to FTEs (0.39) (Figure 3B, right). Additionally, HGSOCs exhibited greater variation within the group (SD = 0.069), in contrast to the FTEs (SD = 0.023).

**Figure 3.**
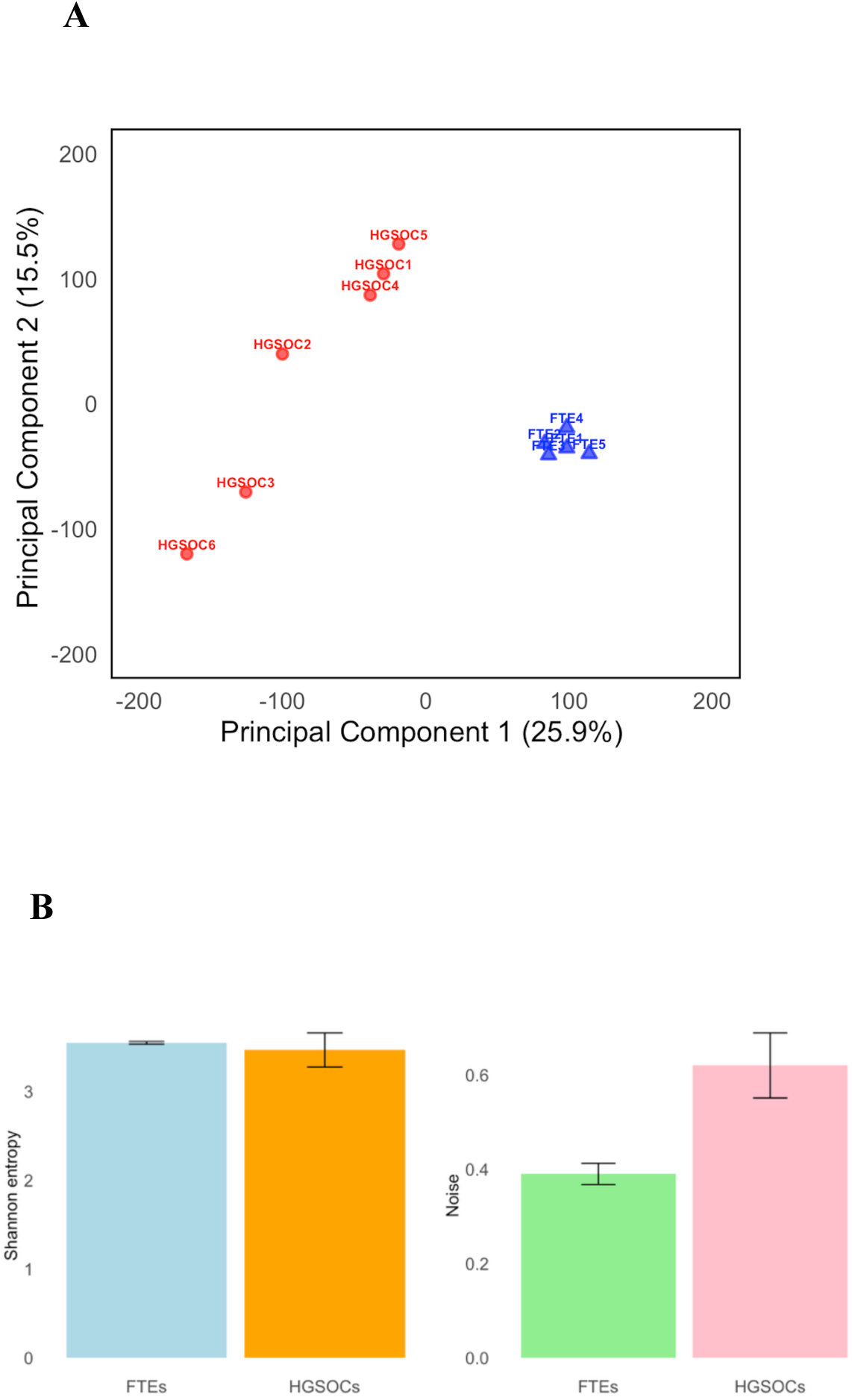
PCA, Shannon Entropy and Noise. (A) PCA plot of the first 2 Principal Components for FTE (red circles) and HGSOC (blue triangles), showing 25.9 variance for the first component and 15.5 variance for the second component. (B) The bar charts of mean Shannon entropy for FTE (blue) and HGSOC (yellow), and mean Noise for FTE (green) and HGSOC (red), and the error bars indicate the variability.

Overall, these analyses allowed us to compare and evaluate the statistical properties of our datasets, and will serve as important tools for assessing the quality of our process-driven transcriptional model that we next develop for synthetic data generation. The goal is for the final model, one for each sample (FTE and HGSOC), to generate synthetic transcriptome-wide expressions that closely mirror the experimental data analyzed here.

### Development of Transcriptional Regulatory (TR) Models for FTEs and HGSOCs

Differential equations have long been a cornerstone of systems biology research, widely used to study the complex dynamic behaviors of biochemical reactions and networks [28,29,30]. The models developed have been successfully applied to investigate various biological processes, including glycolysis, immune and cancer signal transduction, circadian rhythms, and many other essential cellular processes, largely in a deterministic framework [31,32,33,34].

Transcriptional processes are inherently noisy, which is primarily attributed to both intrinsic and extrinsic factors [12,35]. Intrinsic noise typically arises from the low copy number of molecules and the probabilistic or stochastic nature of molecular events, while extrinsic noise is often caused by factors such as fluctuations in molecular expressions between cells, environmental variations, or technical inconsistencies [36,37]. To model intrinsic molecular events, stochastic algorithms based on biochemical reactions with differential equations have been widely adopted [38,39]. For instance, we previously developed a stochastic transcriptional model to examine the variations in transcriptome-wide gene expression noise in developmental cells along their lineage [17].

Extrinsic noise, on the other hand, can be represented by Gaussian or white noise [12,40], often added to stochastic events to model and analyze the total noise in a biochemical system [41]. Thus, here we adopted a stochastic modeling approach based on Gillespie algorithm (intrinsic) with additive Gaussian (extrinsic) noise to investigate the transcriptome-wide expressions of FTE and HGSOC samples (Figure 4A). The rate equations governing gene expression of the *i^th^* gene, *G_i_*, is defined by 3 differential equations:

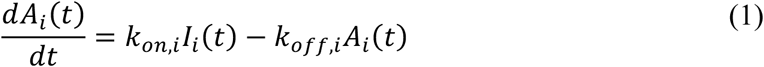

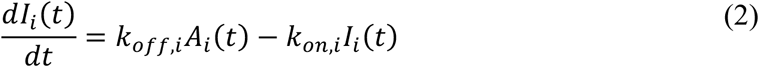

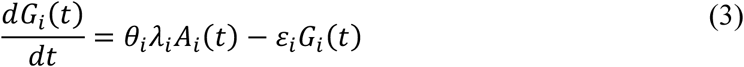

**Figure 4.**
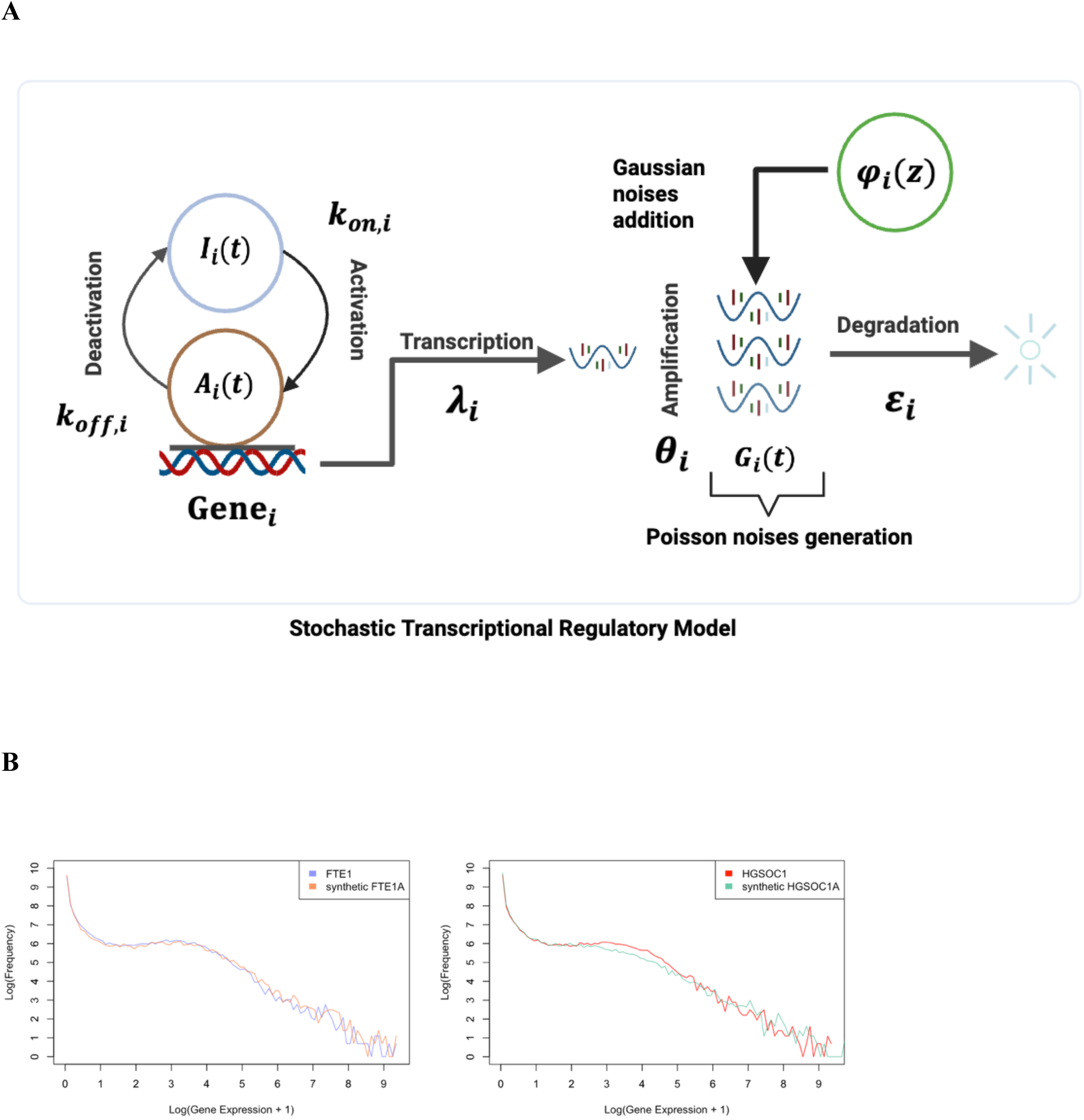
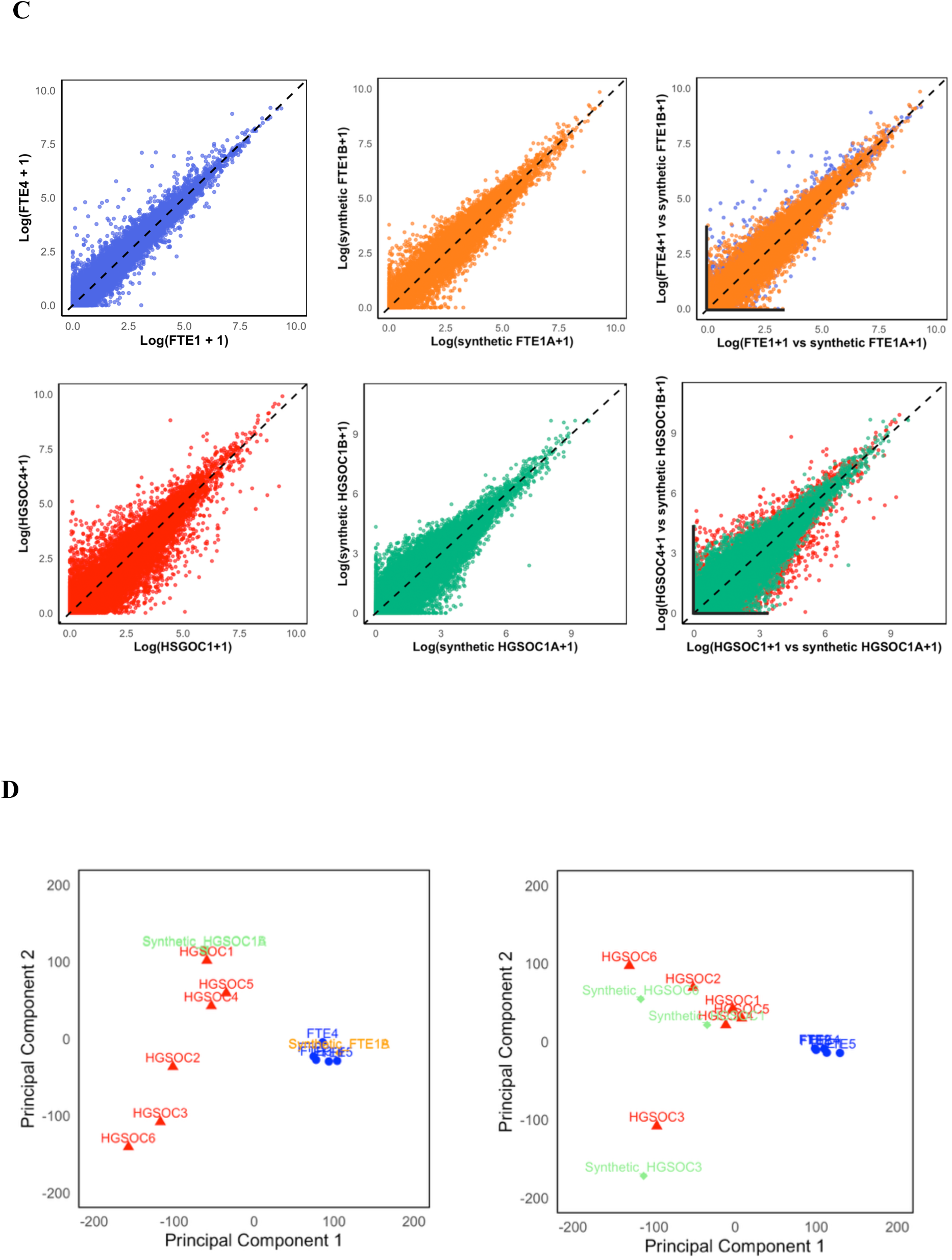
Stochastic gene transcriptional regulatory (TR) model and synthetic data generation. (A) Schematic of the TR model. The intrinsic (Poisson) noises are generated by Gillipse algorithm and the extrinsic (Gaussian) noises are integrated together with varying the 5 parameters to fit the model simulation with actual gene expressions. (B) Model generated synthetic FTEs (orange, dotted) versus experimental FTEs (blue) gene expression distributions (top), and synthetic HGSOCs (green, dotted) versus experimental HGSOCs (red) gene expression distributions (bottom). (C) Scatter plots of experimental FTEs (blue, top left), synthetic FTEs (orange, top middle), synthetic FTEs versus experimental FTEs (top left), and scatter plots of experimental HGSOCs (red, bottom left), synthetic HGSOCs (green, bottom middle), and synthetic HGSOCs versus experimental HGSOCs (bottom left). (D) Left: PCA plot of 2 synthetic FTE1s (yellow rhombus) and 2 HGSOC1s (green square). Right: The revised PCA plot of synthetic HGSOC1,3&6 (green square) with 2400 PC2 loading genes removed.

where the promoter activation of each gene is defined by active (*A_i_*) and inactive (*I_i_*) states (2-state model), and the transition rates between the states are defined by two parameters, *k_on,i_* and *k_off,i_*. The model constitutes of 39,761 gene ‘units’, where each gene dynamics is governed by 5 kinetic parameters [17,42,43]; transcription rate (*λ*), degradation rate (*ε*), promoter activation (*k_on_*), deactivation (*k_off_*) rate constants and the transcriptional amplification process [44], i.e. number of transcripts produced per activation event, is controlled by *θ*.

The final *G_fi_* is superposed with Gaussian noise, 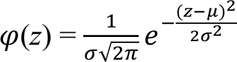

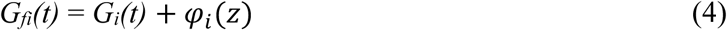

where *z* represents the grey level, *μ* the mean grey value, and *σ* its standard deviation (Methods). By controlling the 5 parameters for each gene, we simulate the gene expressions for all genes for all samples. During the simulations, three parameter values (*k_on_*, *k_off_* and *θ*) were varied or searched for each condition. For transcription (*λ*) and degradation (*ε*) rates, we used the statistical distributions of available experimental data [45,46,47,48], where the degradation rates followed a lognormal distribution and the transcription rates are represented by *T_i_*(*k_on,i_* + *k_off,i_*)/*θ_i_k_on,i_*, where the *T_i_* are time-averaged RNA transcription rates following a lognormal distribution as well.

### Generating Synthetic Transcriptome-wide Expressions for FTEs and HGSOCs

Using the description above, we simulated our stochastic transcriptional regulatory (TR) model to fit both FTEs and HGSOCs transcriptome-wide expressions across all 11 available samples, adjusting all five model parameters for each gene. We assumed that both the transcription rate (*λ*) and degradation rate (*ε*) for each gene follow lognormal distributions, with the mean (*μ*) and standard deviation (*σ*) estimated for our gene expression dataset (n = 39,761 genes) [17].

For our Gaussian noises, we tried several values of *μ* and *σ* for all samples as an initial estimate. However, the stochastic noises with additive Gaussian noises revealed unsatisfactory outcomes (Fig. S3). Notably, several recent studies investigating biological noise in signal transduction, bistability, and cellular heterogeneity have indicated that multiplicative noise provides a more realistic representation. Therefore, we next investigated a Gaussian noise component that was combined multiplicatively with stochastic noise to generate the final *G_fi_* as [49,50,51,52]:

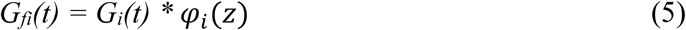

We determined *μ* = 1.3 and *σ* = 0.35 for Gaussian noises generally fitted to all all experimental data. For FTE, we obtained *μ* = 0.57 and *σ* = 2.41 for *λ* using Zipf’s law with *r* = 0.794, and *μ* = −1.05 and *σ* = 0.78 for *ε*. By randomly sampling *λ_i_* and *ε_i_* from the lognormal distribution, and selecting *k_on,i_*, *k_off,i_*, and *θ_i_* to match the actual expression values for the *i_th_* gene using a genetic algorithm, we determined fixed values of *k_on,i_* = 90, *k_off,i_* = 2, and *θ_i_* = 5000 for all genes (Methods). That is, by fixing *k_on,i_*, *k_off,i_*, and *θ_i_* and randomly sampling *λ_i_* and *ε_i_* from the lognormal distributions for all 39,761 genes, we were able to simulate FTE transcriptomes (FTE1 to FTE5) that closely recapitulate the actual gene expression distributions (Figure 4B, left, and Fig. S4). It is important to note that the five FTE simulations used the same model parameters, with differences arising only from the stochastic variations of the model simulations for each run.

To generate synthetic HGSOC expressions, the transcription rate (*λ*) for each gene also follows lognormal distributions, with *μ* = 0.37 and *σ* = 2.05, as determined by fitting Zipf’s law with an exponent *r* = 0.793 to their expression dataset and degradation rate (*ε*) follows lognormal distributions, with *μ* = −0.44 and *σ* = 0.92. Additionally, we set the values of *k_on,i_* = 85, *k_off,i_* = 30, and *θ_i_* = 4000. The HGSOC distributions (HGSOC1 to HGSOC6) also seem to well recapitulate the actual gene expression distributions (Figure 4B, right, and Fig. S4).

To assess the reliability and accuracy of the synthetic FTEs and HGSOCs generated by our TR model, we performed various statistical and data analytic comparisons with the experimental data (Figure 4C, Table 2). The comparisons include pair-wise scatter plots of real (Blue for FTEs & Red for HGSOCs) versus synthetic (Orange for FTEs & Green for HGSOCs) data (Figure 4C, Fig. S4). We assessed the transcriptome-wide Pearson Correlation, Shannon entropy, and noise, comparing real data with synthetic data generated using both stochastic and Gaussian noises (Table 2). Our overall results show that the model and real expressions’ similarity ranges between 82 to 99% across all metrics investigated.

**Table 2.** Similarity between experimental and synthetic transcriptomes of FTEs and HGSOCs using Noise, Pearson correlation coefficient, and Shannon entropy.

| FTE |  | Experiments | Synthetics | Similarity |
| --- | --- | --- | --- | --- |
| F1 vs F4 | Noise | 0.428 | 0.442 | 96.8% |
|  | PCC | 0.953 | 0.942 | 98.8% |
|  | Shannon entropy | 3.555 | 3.477 | 97.8% |
| F2 vs F4 | Noise | 0.422 | 0.457 | 92.3% |
|  | PCC | 0.934 | 0.958 | 97.5% |
|  | Shannon entropy | 3.521 | 3.401 | 96.6% |
| F3 vs F4 | Noise | 0.423 | 0.388 | 91.7% |
|  | PCC | 0.933 | 0.940 | 99.3% |
|  | Shannon entropy | 3.515 | 3.382 | 96.2% |
| F4 vs F1 | Noise | 0.428 | 0.475 | 90.1% |
|  | PCC | 0.953 | 0.941 | 98.7% |
|  | Shannon entropy | 3.535 | 3.435 | 97.2% |
| F5 vs F4 | Noise | 0.416 | 0.368 | 88.5% |
|  | PCC | 0.906 | 0.922 | 98.3% |
|  | Shannon entropy | 3.537 | 3.451 | 97.6% |

| HGSOC |  | Experiments | Synthetics | Similarity |
| --- | --- | --- | --- | --- |
| H1 vs H4 | Noise | 0.502 | 0.490 | 97.6% |
|  | PCC | 0.860 | 0.909 | 94.6% |
|  | Shannon entropy | 3.473 | 3.249 | 93.6% |
| H2 vs H4 | Noise | 0.570 | 0.539 | 94.6% |
|  | PCC | 0.895 | 0.950 | 94.2% |
|  | Shannon entropy | 3.382 | 3.221 | 95.2% |
| H3 vs H4 | Noise | 0.643 | 0.572 | 89.0% |
|  | PCC | 0.887 | 0.969 | 91.5% |
|  | Shannon entropy | 3.304 | 3.067 | 92.8% |
| H4 vs H1 | Noise | 0.502 | 0.510 | 98.4% |
|  | PCC | 0.860 | 0.974 | 88.3% |
|  | Shannon entropy | 3.373 | 3.081 | 91.3% |
| H5 vs H1 | Noise | 0.526 | 0.607 | 86.7% |
|  | PCC | 0.721 | 0.878 | 82.1% |
|  | Shannon entropy | 3.550 | 3.260 | 91.8% |
| H6 vs H4 | Noise | 0.668 | 0.602 | 90.1% |
|  | PCC | 0.803 | 0.954 | 84.2% |
|  | Shannon entropy | 3.024 | 2.789 | 92.2% |

We revisited the PCA plot of FTEs and HGSOCs along the first two components to assess the similarity between the synthetic and real data (Figure 4D, left). For FTEs, the synthetic data (for 2 simulations) appears to align reasonably well. However, for HGSOCs, only HGSOC1 is adequately replicated (for 2 simulations). The generally high gene expression variability between the HGSOC samples complicates the model’s ability to generate highly variable gene expressions using fixed parameters and relying solely on stochastic and Gaussian noises. Therefore, it is essential to identify the genes responsible for the significant variability between HGSOCs, particularly on the PC2 axis.

We hypothesized that certain genes contribute to the high variability observed in PC2 values among HGSOC samples. To investigate this, we identified genes with high PC2 loadings across all HGSOCs and systematically removed them in sets of the top 100 loading genes. After excluding the top 2,400 PC2-loading genes, the HGSOC samples organized into three relatively distinct clusters (Fig. S5). We then used this reduced transcriptome to fit the HGSOC model to the three experimental clusters (Figure 4D, right), resulting in one model per cluster. Notably, if all genes were retained, the diversity along the PC2 axis would necessitate fitting each sample individually, requiring six models for the HGSOCs.

Finally, we identified the L-shaped toggle genes in the finalized synthetic data and compared them with their real counterparts, in which 4,833 and 5,232 toggle genes were identified from synthetic FTE and HGSOC simulations, respectively (Figure 4C, L-shaped black lines). The number of toggle genes generated from our model indicated a remarkable similarity (around 80%) with the number of experimental toggle genes. This demonstrates that our TR model, consisting of three differential equations and five model parameters, can be fine-tuned to accurately match transcriptome-wide expressions for both FTE and HGSOC.

### DE analysis between FTEs and HGSOCs

To identify differentially expressed (DE) genes between FTEs and HGSOCs, we employed Limma package in R, using the Benjamini-Hochberg correction method (for a 2-fold change and *p* < 0.05) [53]. A total of 2,123 DE genes were identified, and the gene enrichment analysis, using *clusterProfiler* in R [54], revealed that the most significant biological pathways (based on gene counts and *p*-values) are related to cellular structure and functions, including microtubule-based movement and cilium organization (Figure 5A). We also conducted gene enrichment analyses for the 2,400 variable high PC2 loading genes, mentioned above, to identify the biological processes associated with the variability between the HGSOCs (Figure 5B). Our analysis revealed that the enriched functions were closely related to broad biological processes, such as vesicle organization and Golgi vesicle transport.

**Figure 5.**
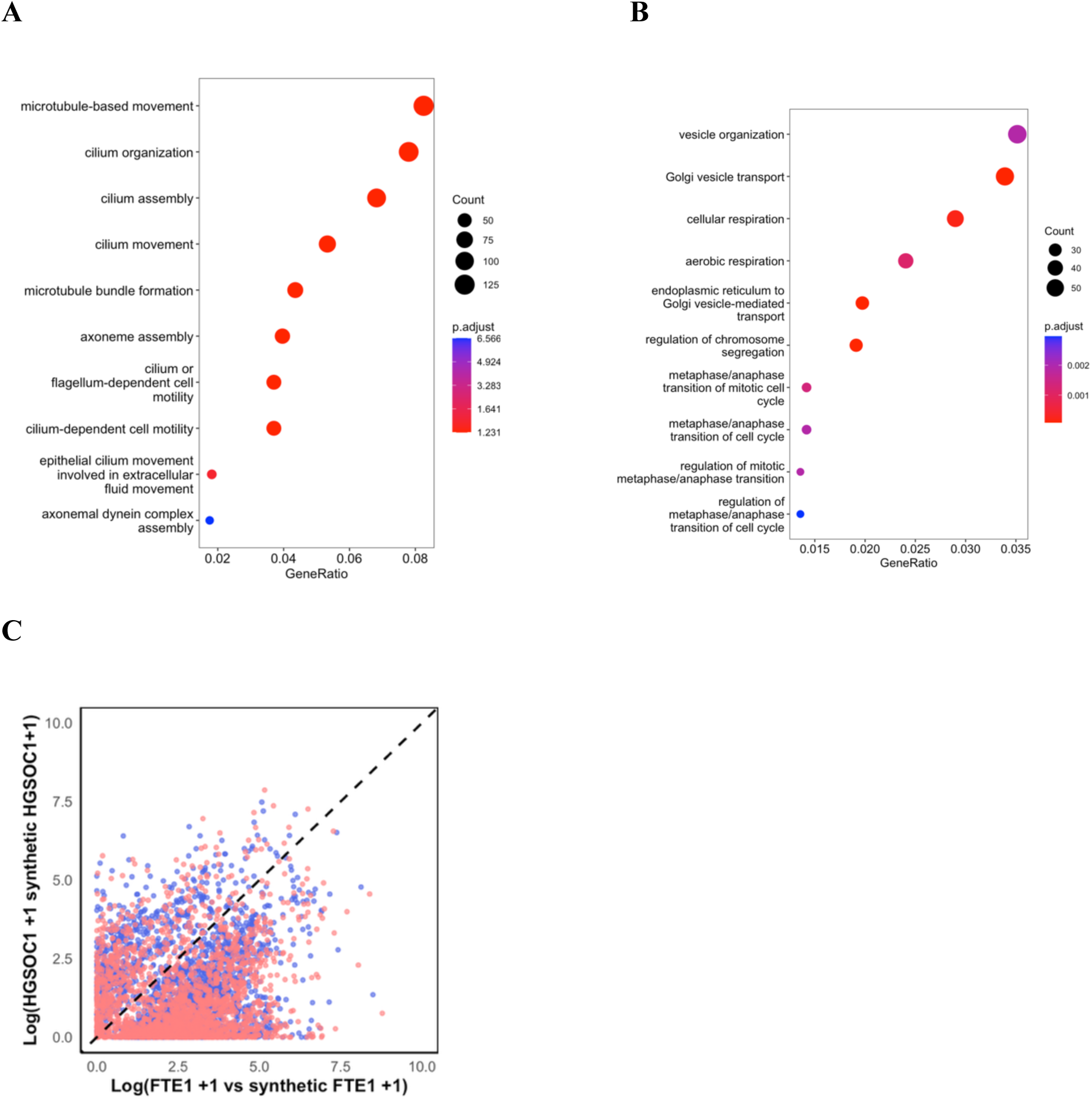
Gene Enrichment Analysis of DE and PC2 loading genes. (A) The dot plots show the top significant biological process of DE genes. The size of dots indicates the number of genes associated with the GO term, and the colour of dots indicates the *p*-adjusted values. (B) The dot plots show the top significant biological process of PC2 loading genes. The size of dots indicates the number of genes associated with the GO term, and the colour of dots indicates the *p*-adjusted values. (C) Scatter plot of experimental FTE1 and HGSOC1 (blue) versus synthetic FTE1 and HGSOC1 (red) for all 2,123 DE genes.

Next, we used our TR model to uncover the regulatory properties of the top 100 DE genes, selected based on their adjusted *p*-value. To accomplish this, we first mapped the actual genes to their synthetic counterparts using scatter plots (Figure 5C). The scatter plots showed the plots of all DE genes (blue) and their synthetic counterparts (red). Although we could map all real DE genes with the synthetic data for revealing their model parameters, for illustration, we focused only on the top 100 DE genes. After matching, we then determined the *k_on_*, *k_off_*, amplification factor *θ*, transcriptional rate *λ*, and the degradation rate *ε* for each of the top 100 DE genes (Table S1). We then performed t-tests to compare the transcriptional and degradation rate parameters between FTEs and HGSOCs for the top 10 differentially expressed (DE) genes. The analysis revealed *p*-values below the 0.05 threshold for both transcriptional and degradation rates, indicating significant differences between FTEs and HGSOCs (Table S1). This approach allows the TR model to provide a mechanistic insight into how each of these DE genes is regulated in control versus disease samples It is important to note that certain model parameters were similar between the control and disease groups, whereas others differed significantly. These patterns may reflect underlying co-regulatory relationships among genes that are either shared or distinct between the two conditions. [55][56].

For instance, in Table 2, while *k_on_*, *θ*, and *ε* are relatively similar between FTEs and HGSOCs, *k_off_* is notably lower in FTEs (∼2) compared to HGSOCs (∼30), and *λ* is significantly higher in FTEs than in HGSOCs, especially for the highly variable genes. Thus, when looking at the top 10 DE genes, these analyses suggest that they exhibit greater variability in transcription factor binding, indicating an increased likelihood of transcriptional bursting and elevated activation levels compared to variations observed in degradation mechanisms, such as those mediated by miRNAs or other ncRNAs, consistent with previous reports [57,58].

We further investigated the functions of the top 10 DE genes and identified that these genes were all protein-coding genes. Genes like *GSTA3* (detoxification) and *ADH1B* (alcohol metabolism, tied to fetal alcohol syndrome) protect cells from damage from toxic exogenous and metabolize ethanol [59,60]. *SNTN* and *DNAI2* are critical for ciliary function, with mutations causing primary ciliary dyskinesia [61,62]. C6 contributes to immune defence by forming the membrane attack complex, and *KIF19* (a microtubule motor) may enable ATO hydrolysis activity and is associated with hydrops fetalis [63,64]. Moreover, *ANKRD66* may regulate protein modifications via NAD+ [65].

We next explored the relationships of enriched terms between the highly PC2 loading and DE genes (Figure 6A, left). There are 4,993 terms that are common between them and several enriched functions are displayed (Figure 6A, right). Moreover, we used Cystoscope to generate the DE, PC2 loading, and random gene networks. The parameters of color and size were varied to demonstrate the connectivity of each node as the bigger size and brighter color indicate a higher number of interactions of the node. We identified DE genes and PC2 loading genes networks exhibited a significantly larger number of connected nodes in the centre, compared to the random gene network, which only revealed a few highly connected nodes (Figure 6B). Lastly, we plotted the density plot for the 3 networks, and observed only the random genes network follows a power-law (Figure 6C) [66]. This is conceivable as DE and high PC2 loading genes are specifically associated with cancer and thus are not expected to show the whole or random network behaviour [27].

**Figure 6.**
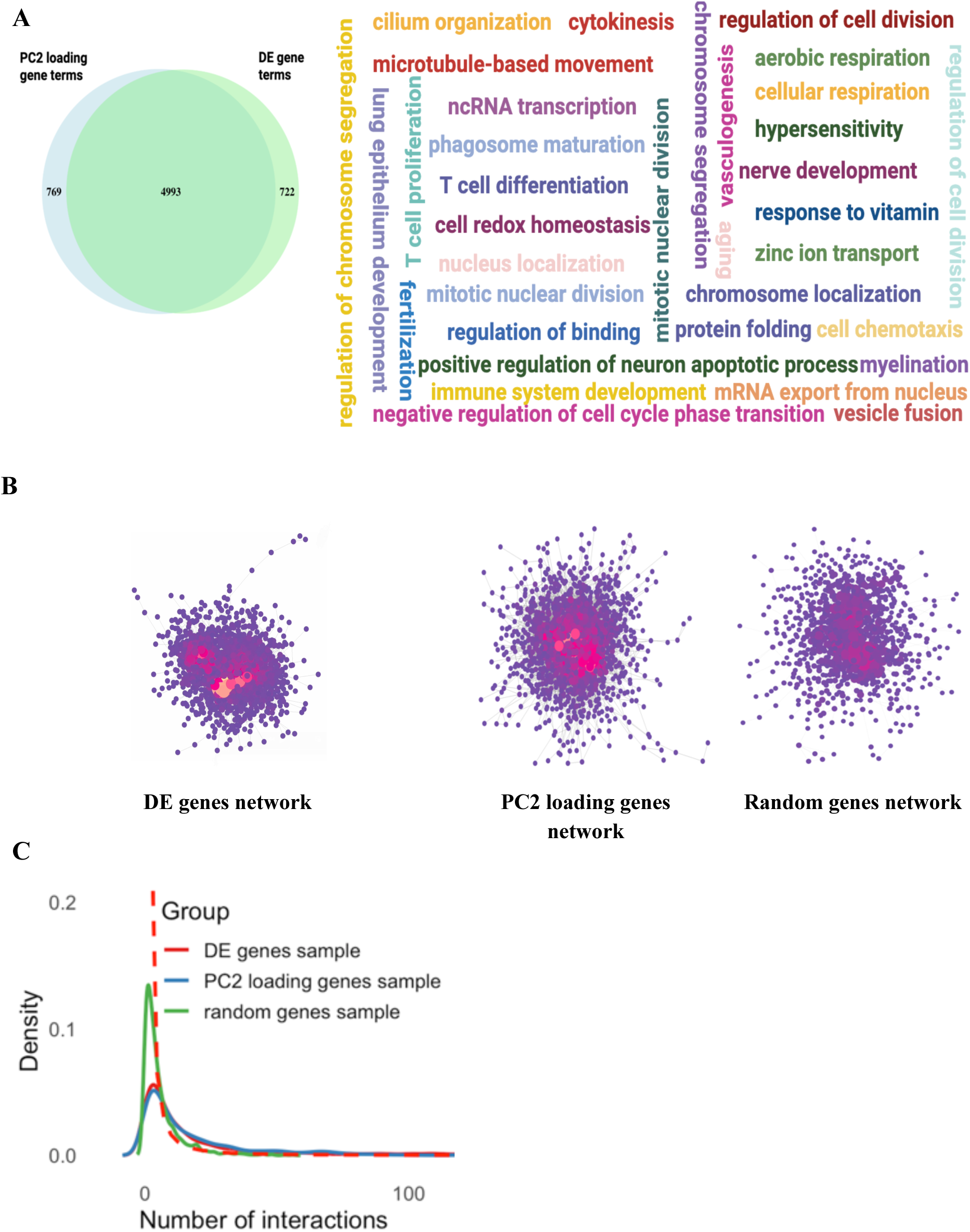
Common Gene Enrichments between DE and PC2 loading genes and Networks. (A) Venn Diagram showing a significant number of enriched terms from the terms that PC2 loading genes (blue) are associated closely with DE genes (green). (B) Word cloud figure visually represents various shared biological processes and terms between PC2 loading gene terms and DE gene terms. (C) GO networks for DE (left), PC2 loading (central), and randomly sampled genes (right) generated from Cystoscope. The random genes have the same number as the number of PC2 loading genes, and the nodes in networks are scaled to a yellower colour and a bigger size. (D) The density plot of the number of interactions of DE (red line), PC2 loading (blue line), and randomly sampled genes (green line), and the power law distribution is represented by a red dashed line.

### Exploring the universality of TR modelling approach using an independent dataset on cardiomyocytes and single cell RNA-seq of ovarian cancer

To assess the general applicability of our approach, we analyzed a cardiomyocyte dataset under formin Diaphanous (DIAPH) and scramble (Scr) knockdown conditions (GSE214149) [67]. For biological replicates under DIAPH knockdown, we obtained *µ* = 0.21 and *σ* = 2.07 for *λ*, and *µ* = −0.58 and *σ* = 0.88 for *ε*, with fixed values of *k*_on_ = 10, *k*_off_ = 7.5, and *θ_i_* = 5000. For biological replicates under Scr knockdown, we obtained *µ* = 0.18 and *σ* = 2.05 for *λ*, and *µ* = −0.48 and *σ* = 0.93 for *ε*, with fixed values of *k*_on_ = 9, *k*_off_ = 12.5, and *θ_i_* = 4500. Using these parameters, we successfully generated synthetic data that recapitulated the biological patterns observed in the real replicates (Figure S6, Table 4).

We also analyzed expression distributions, transcriptome-wide scatter plots, noise, Shannon entropy, and Pearson’s correlation coefficients (Figure S6 and Table 4). The results demonstrated a high degree of similarity between the real and synthetic replicates, confirming that our TR model can reproduce the statistical properties of real transcriptomes. This similarity is further evident in the expression distributions and transcriptome-wide scatter plots, where the real and synthetic data largely overlap (Fig. S6).

**Table 3.**
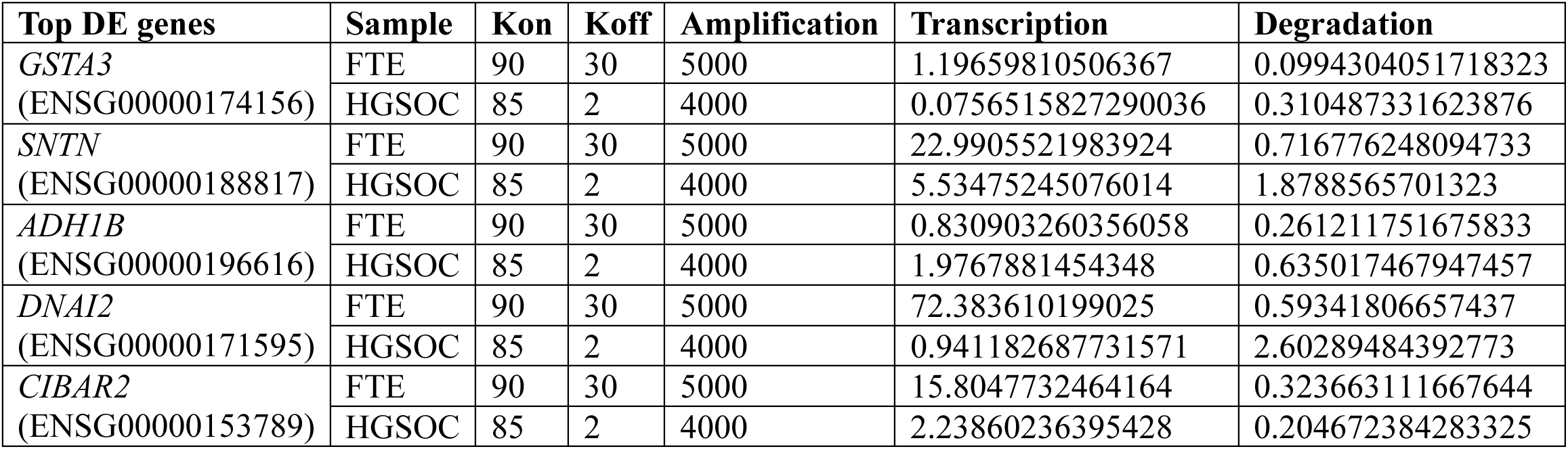
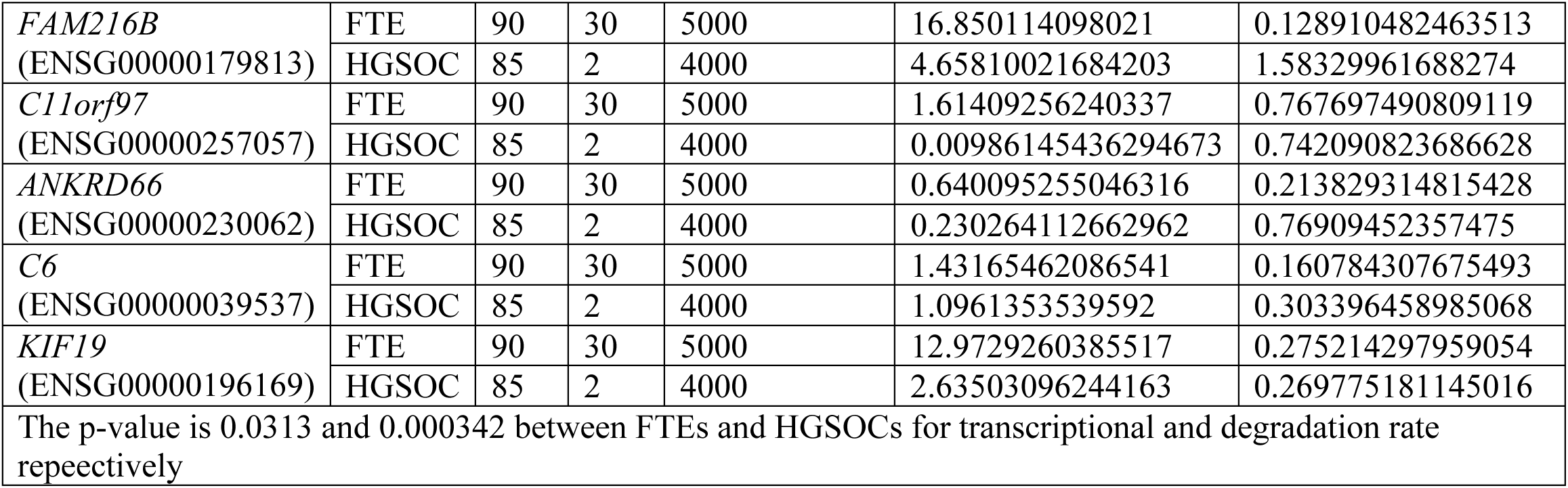
Top 10 DE genes and their TR model parameter values.

**Table 4.** Similarity between experimental and synthetic transcriptomes of DIAPHs and Scrs using Noise, Pearson correlation coefficient, and Shannon entropy.

| DIAPH | Experiments<br>DIAPH2vsDIAPH3 | Synthetic<br>(Poisson) | Similarity |
| --- | --- | --- | --- |
| Noise<br>Level | 0.0427 | 0.0406 | 95.1% |
| PCC | 0.999 | 0.999 | 100% |
| Shannon<br>entropy | 3.474 | 3.412 | 98.2% |

| Scr | Experiments<br>Scr4vsScr1 | Synthetic<br>(Poisson) | Similarity |
| --- | --- | --- | --- |
| Noise<br>Level | 0.0573 | 0.0565 | 98.6% |
| PCC | 0.999 | 0.999 | 100% |
| Shannon<br>entropy | 3.371 | 3.344 | 99.2% |

Finally, we applied our stochastic transcriptional model to an ovarian single-cell RNA sequencing dataset (GSE192898) [68]to evaluate its performance on single-cell transcriptomic data. By adjusting *µ* = 0.27 and *σ* = 1.95 for *λ*, *µ* = −0.45 and *σ* = 1.38 for *ε*, and setting *k*_on_= 0.9, *k*_off_ = 10, and *θ_i_* = 5000, the model successfully simulated the top 2000 genes for single-cell transcriptomes. Statistical analyses, expression distributions, and transcriptome-wide scatter plots were then used to assess its accuracy and reliability, demonstrating a high concordance between real and synthetic data (Figure S7, Table 5).

**Table 5.**
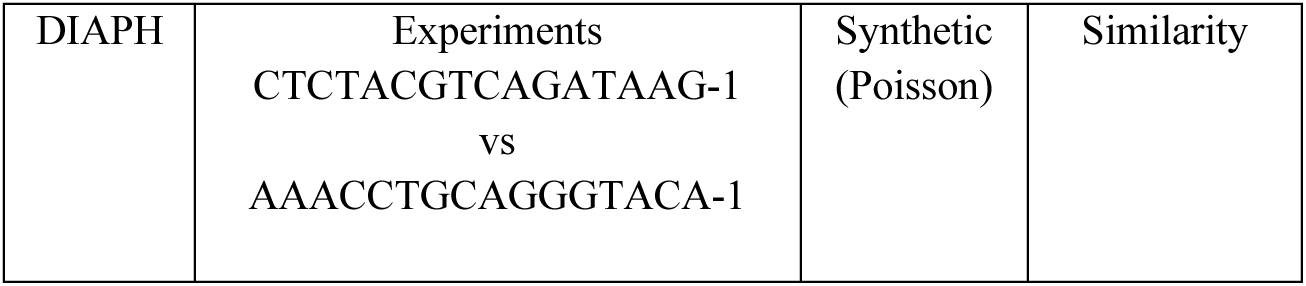
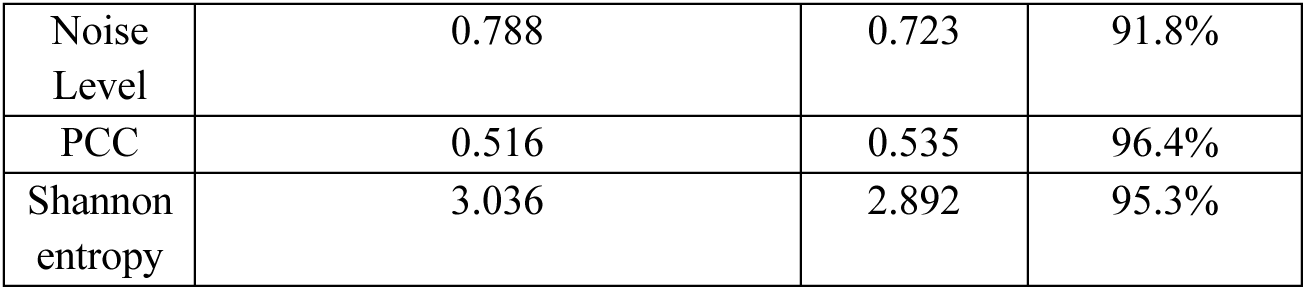
Similarity between experimental and synthetic Single cell RNA-Sequening data using Noise, Pearson correlation coefficient, and Shannon entropy.

Overall, we have demonstrated that our TR model simulations can be directly compared to the experimental data for all samples, achieving a high level of accuracy across several metrics, including expression levels, correlations, noise, Shannon entropy, and PCA scores. Using this model, we were able to identify the regulatory parameters for key DE genes.

## Discussion

Studies on differential gene expressions between disease and normal cells have offered valuable insights into identifying candidate biomarkers and proposing new drug targets [69]. However, these studies often fall short of providing a comprehensive understanding of transcriptome-wide expression regulation. In this study, we introduce a stochastic transcriptional regulatory (TR) model designed to examine gene regulation of differentially expressed genes between control and ovarian samples from a transcriptome-wide perspective.

Initially, we generated expression distributions and transcriptome-wide scatter plots for all FTE and HGSOC replicates. By incorporating statistical metrics such as Pearson Correlation Coefficient (PCC), Shannon entropy, and noise, we observed that an increase in gene expression variability exists from FTE to HGSOC replicates. This is expected, as cancer cells are known to exhibit greater variability compared to normal cells from the same tissue.

Next, we introduced our stochastic TR model, which includes five regulatory parameters. By adjusting these parameters to follow specific known statistical distributions or constant experimentally fitted values, we successfully generated various synthetic datasets for each FTE and HGSOC sample. Comparing the expression distributions and scatter plots of the synthetic FTE and HGSOC replicates with experimental counterpart data demonstrated a good fit, verifying the reliability of our model. Further analysis of transcriptome-wide noise, PCC, and Shannon entropy metrics all showed comparable results, between 82-99% accuracy.

Additionally, we performed PCA on the datasets and observed a clear separation between FTE and HGSOC replicates. When we added synthetic FTE and HGSOC data to the PCA plot, we found that the synthetic FTE data clustered closely with their corresponding experimental data. However, for the HGSOC replicates, it was challenging to generate synthetic data that fully captured the variability observed along the PC2 axis, mainly due to the high inherent variability within the replicates, as is typical with cancer cells. This variability highlighted significant tumor heterogeneity, with the emergence of subclusters indicating different molecular states within the tumors. This variability could be attributed to genetic mutations, epigenetic modifications, or interactions within the tumor microenvironment (TME) or the diversity of cells in the TME [70,71,72,73]. To address this, we applied a method to re-cluster the HGSOC samples by removing the high PC2-loading genes. As a result, when we removed 2,400 high PC2-loading genes, the formation of three distinct HGSOC clusters were achieved, for which we only need 1 model for all HGSOCs that could form 3 distinct clusters We then conducted Gene Enrichment Analysis to investigate biological processes, where all DE genes were found to be enriched in functions like microtubule-based movement, cilium organization and axoneme assembly. Next, we analyzed the 2,400 high-PC2 loading genes, which revealed significant enrichment in functions related to vesicle organization and Golgi vesicle transport. Nevertheless, we also noticed a significant commonality in the enriched functions between DE genes and PC2-loading genes. Notably, when tested on power-law distributions, we observed that random networks the size of PC2 loading genes display power-law but not DE and PC2 loading genes, which possessed a larger number of highly connected nodes.

We also examined the parameter values for the top 100 DE genes in the synthetic data generated TR model and found the high variability of DE genes are mainly due to quantal, higher transcriptional, and higher degradation rates for HGSOC compared with FTEs, leaving to higher expression diversity between them. This information, in the future, may be used to control specific ovarian cancer biomarkers’ transcriptional expressions *in silico* before new regulatory targets may be developed to test them *in vivo*, to better regulate ovarian cancer growth and proliferations.

In conclusion, our analysis, which involved mapping synthetic gene expression data generated by the TR model to the real counterparts, demonstrated its reliability and accuracy. This enabled us to uncover the transcriptional regulatory mechanisms for DE genes between control and ovarian cancer cells. Our modelling approach can be applied to any gene expression dataset that possesses both control samples with other diseases, where we also tested and verified with another independent dataset on cardiomyocytes.

Thus, our study offers valuable insights into previously unexplored transcriptome-wide expression regulation, which could improve our understanding of transcriptional regulation and pave the way for future RNA-level therapeutic treatments by specifically targeting genes that are critically regulated in disease processes.

## Methods

### Bulk RNA-seq dataset

For high-grade serous ovarian carcinoma (GSE190688), bulk RNA-seq datasets were obtained from the Gene Expression Omnibus (GEO) database using previously published data [18]. The datasets were selected based on the availability of associated healthy patient tissue and tumour samples.

### Data Availability

The Synthetic TR Model source code with user instructions and ovarian cancer dataset can be found on URL: https://github.com/OrcsCheung/Stochastic-Transcriptional-Regulatory-Model.git

### Distribution fitting

In order to determine statistical distribution fitting of each sample in bulk RNA-seq datasets, we used the distribution with lowest Akaike information criterion value to fit for the samples by using *fitdistplus* [74] and to determine parameter estimation from several distributions: log-normal and gamma distribution.

### Pearson correlation coefficient

A Pearson correlation coefficient is a linear correlation coefficient that defines the degree of the relation between two variables. The measure is denoted by *r* and calculated as the covariance of the two variables divided by the product of their standard deviations. The formula of the Pearson correlation coefficient is below:

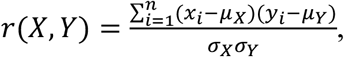

Where *x_i_* and *y_i_* are the expression value for the ith gene in the dataset for each of the two samples and *µ_X_* and *µ_Y_* represent the mean expression of each sample and *σ_X_* and *σ_Y_* represent the standard deviation of the expression values in each sample. The *cor* function of the R *stats* package is used to estimate the correlations [75,76,77].

### Average transcriptome noise calculation

To compute the average transcriptome noise, we need to calculate the squared coefficients of variation of each gene (*i* = 1, …, *m*) between pairs of samples (*j*, *k* = 1, …, *n*) :

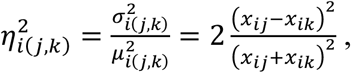

where the *x_ij_* and *x_ik_* represent the expression value of *i^th^* gene in *j^th^* sample and expression value of *i^th^* gene in *k^th^* sample. Then, we need to determine the average noise of each gene for all samples:

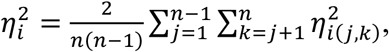

Eventually, we can calculate the average transcriptome noise based on the average noise of each gene for all samples:

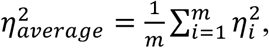

Where the *m* is the total number of genes.

### Gene expression entropy

The entropy value of gene expression for each sample is defined as:

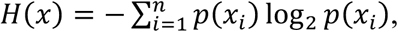

Where *n* is the number of transcriptomes and *p*(*x_i_*) is the probability of gene expression value *x* = *x_i_*. The *entropy.empirical* function of R *entropy* package is used to estimate the correlations [78].

### Toggle gene identification

Toggle genes which exhibit a 0 expression in one sample and a positive expression in another sample between 2 samples in the same condition are defined by Giuliani et al [26]:

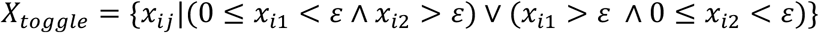

where, *x_ij_* denoted the expression vector of the *i*^th^ gene of any two samples *j=*1,2 in the same conditions. The *ε* is a minimum expression threshold and determined from statistical distribution fitting of gene expression by using the method (Distribution fitting) aforementioned.

### GO analysis

For gene enrichment analysis, only the biological process was performed by using the *clusterProfiler* in R and a threshold of *p-value* <0.05 [54]. Then, the dot plot was used to visualize several top-enriched terms.

### Cystoscope network

StringApp in Cystoscope was used to create networks for DE, PC2 loading and randomly sampled genes [79,80,81]. The networks were examined using the software’s Analyzer, which also calculated the degree parameters for every gene in every network. Degree, or the number of connections per node, and connectivity, or the number of connected components in the subset of nodes, were the parameters taken into consideration. A higher value denotes a network with better connectivity, while a lower value denotes elements that are more disjoint and have poorer connectivity [81].

## Acknowledgement

The authors thank ASTAR Bioinformatics Institute for supporting this work under the core research budget of KS and the Nanyang Technological University for the MSc in Biomedical Data Science Programme of KS and SZ.

## Author Contributions

Z.S. performed the model development, analysis and wrote the paper. K.S conceptualized, supervised, wrote and edited the paper.

## References

[1] Long, E., Wan, P., Chen, Q., Lu, Z. & Choi, J. From function to translation: decoding genetic susceptibility to human diseases via artificial intelligence. Cell Genom. 3, 100320 (2023). DOI: 10.1016/j.xgen.2023.100320

[2] Xu, J., Yang, P., Xue, S., Sharma, B., Sanchez-Martin, M., Wang, F. et al. Translating cancer genomics into precision medicine with artificial intelligence: applications, challenges and future perspectives. Hum. Genet. 138, 109–124 (2019). DOI: 10.1007/s00439-019-01970-5

[3] Yu, J. S. & Bagheri, N. Multi-class and multi-scale models of complex biological phenomena. Curr. Opin. Biotechnol. 39, 167–173 (2016). DOI: 10.1016/j.copbio.2016.04.002

[4] Ruan, D., Young, A. & Montana, G. Differential analysis of biological networks. BMC Bioinformatics 16, 327 (2015). DOI: 10.1186/s12859-015-0735-5

[5] Tu, J.-J., Ou-Yang, L., Zhu, Y., Yan, H., Qin, H. & Zhang, X.-F. Differential network analysis by simultaneously considering changes in gene interactions and gene expression. Bioinformatics 37, 4414–4423 (2021). DOI: 10.1093/bioinformatics/btab502

[6] Grimes, T., Potter, S. S. & Datta, S. Integrating gene regulatory pathways into differential network analysis of gene expression data. Sci. Rep. 9, 5479 (2019). DOI: 10.1038/s41598-019-41918-3

[7] van Dam, S., Võsa, U., van der Graaf, A., Franke, L. & de Magalhães, J. P. Gene co-expression analysis for functional classification and gene-disease predictions. Brief. Bioinform. 19, 575–592 (2018). DOI: 10.1093/bib/bbw139

[8] Xiao, Z., Dai, Z. & Locasale, J. W. Metabolic landscape of the tumor microenvironment at single cell resolution. Nat. Commun. 10, 3763 (2019). DOI: 10.1038/s41467-019-11738-0

[9] Wu, F., Fan, J., He, Y. et al. Single-cell profiling of tumor heterogeneity and the microenvironment in advanced non-small cell lung cancer. Nat. Commun. 12, 2540 (2021). DOI: 10.1038/s41467-021-22801-0

[10] Liu, Z. L., Meng, X. Y., Bao, R. J. et al. Single cell deciphering of progression trajectories of the tumor ecosystem in head and neck cancer. Nat. Commun. 15, 2595 (2024). DOI: 10.1038/s41467-024-46912-6

[11] Li, Z. & Peng, G. Spatial transcriptomics: new dimension of understanding biological complexity. Biophys. Rep. 8, 119–135 (2022). DOI: 10.52601/bpr.2021.210037

[12] Swain, P. S., Elowitz, M. B. & Siggia, E. D. Intrinsic and extrinsic contributions to stochasticity in gene expression. Proc. Natl Acad. Sci. USA 99, 12795–12800 (2002). DOI: 10.1073/pnas.162041399

[13] Selvarajoo K. Understanding multimodal biological decisions from single cell and population dynamics. Wiley Interdiscip Rev Syst Biol Med. 2012 Jul-Aug;4(4):385–99. doi: 10.1002/wsbm.1175

[14] Pedraza, J. M. & Paulsson, J. Effects of molecular memory and bursting on fluctuations in gene expression. Science 319, 339–343 (2008). DOI: 10.1126/science.1144331

[15] Paulsson, J., Berg, O. & Ehrenberg, M. Stochastic focusing: fluctuation-enhanced sensitivity of intracellular regulation. Proc. Natl Acad. Sci. USA 97, 7148–7153 (2000). DOI: 10.1073/pnas.110057697

[16] Gorin, G., Vastola, J. J., Fang, M. et al. Interpretable and tractable models of transcriptional noise for the rational design of single-molecule quantification experiments. Nat. Commun. 13, 7620 (2022). DOI: 10.1038/s41467-022-34857-7

[17] Piras, V., Tomita, M. & Selvarajoo, K. Transcriptome-wide variability in single embryonic development cells. Sci. Rep. 4, 7137 (2014). DOI: 10.1038/srep07137

[18] Dou, Z. et al. HJURP promotes malignant progression and mediates sensitivity to cisplatin and WEE1-inhibitor in serous ovarian cancer. Int. J. Biol. Sci. 18, 1188–1210 (2022). DOI: 10.7150/ijbs.65589

[19] Benesty, J., Chen, J., Huang, Y. & Cohen, I. Pearson correlation coefficient. In Noise Reduction in Speech Processing (eds. Benesty, J. et al.) 71–84 (Springer, 2009). DOI: 10.1007/978-3-642-00296-0_5

[20] Hou, J., Ye, X., Feng, W. et al. Distance correlation application to gene co-expression network analysis. BMC Bioinformatics 23, 81 (2022). DOI: 10.1186/s12859-022-04609-x.

[21] de Siqueira Santos, S., Takahashi, D. Y., Nakata, A. & Fujita, A. A comparative study of statistical methods used to identify dependencies between gene expression signals. Brief. Bioinform. 15, 906–918 (2014). DOI: 10.1093/bib/bbt051.

[22] Shannon, C. E. A mathematical theory of communication. Bell Syst. Tech. J. 27, 379–423, 623-656 (1948).

[23] Ameri, A. J., Lewis, Z. A. Shannon entropy as a metric for conditional gene expression in Neurospora crassa. G3 Genes Genomes Genet. 11, jkab055 (2021). DOI: 10.1093/g3journal/jkab055

[24] Bar-Even, A. et al. Noise in protein expression scales with natural protein abundance. Nat. Genet. 38, 636–643 (2006). DOI: 10.1038/ng1807

[25] Giuliani A, Bui TT, Helmy M, Selvarajoo K. Identifying toggle genes from transcriptome-wide scatter: A new perspective for biological regulation. Genomics. 2022 Jan;114(1):215–228.

[26] Sirbu, O., Agarwal, G., Giuliani, A. & Selvarajoo, K. Understanding the role of toggle genes in chronic lymphocytic leukemia proliferation. bioRxiv [Preprint] at 10.1101/2025.03.31.646494 (2025).

[27] Sirbu, O., Helmy, M., Giuliani, A. et al. Globally invariant behavior of oncogenes and random genes at population but not at single cell level. npj Syst. Biol. Appl. 9, 28 (2023). DOI: 10.1038/s41540-023-00290-9

[28] Elowitz, M. & Leibler, S. A synthetic oscillatory network of transcriptional regulators. Nature 403, 335–338 (2000). DOI: 10.1038/35002125

[29] Niu, Y., Burrage, K. & Chen, L. Modelling biochemical reaction systems by stochastic differential equations with reflection. J. Theor. Biol. 396, 90–104 (2016). DOI: 10.1016/j.jtbi.2016.02.010

[30] Fages, F., Gay, S. & Soliman, S. Inferring reaction systems from ordinary differential equations. Theor. Comput. Sci. 599, 64–78 (2015). DOI: 10.1016/j.tcs.2014.07.032

[31] Philipps, M., Schmid, N. & Hasenauer, J. Current state and open problems in universal differential equations for systems biology. npj Syst Biol Appl 11, 101 (2025). DOI: 10.1038/s41540-025-00550-w

[32] Makhlouf AM, El-Shennawy L, Elkaranshawy HA. Mathematical Modelling for the Role of CD4^+^T Cells in Tumor-Immune Interactions. Comput Math Methods Med. 2020 Feb 19;2020:7187602. doi: 10.1155/2020/7187602.

[33] Kim, R., Reed, M.C. A mathematical model of circadian rhythms and dopamine. Theor Biol Med Model 18, 8 (2021). doi:10.1186/s12976-021-00139-w.

[34] Piras V, Hayashi K, Tomita M, Selvarajoo K. Enhancing apoptosis in TRAIL-resistant cancer cells using fundamental response rules. Sci Rep. 1, 144 (2011). doi: 10.1038/srep00144.

[35] Elowitz, M. B., Levine, A. J., Siggia, E. D. & Swain, P. S. Stochastic gene expression in a single cell. Science 297, 1183–1186 (2002). DOI: 10.1126/science.1070919

[36] Ham, L., Coomer, M.A., Öcal, K. et al. A stochastic vs deterministic perspective on the timing of cellular events. Nat Commun 15, 5286 (2024). doi:10.1038/s41467-024-49624-z

[37] Mitchell S, Hoffmann A. Identifying Noise Sources governing cell-to-cell variability. Curr Opin Syst Biol 8, 39–45 (2018). doi: 10.1016/j.coisb.2017.11.013.

[38] Thomas, P., Popović, N. & Grima, R. Phenotypic switching in gene regulatory networks. Proc. Natl Acad. Sci. USA 111, 6994–6999 (2014). DOI: 10.1073/pnas.1400049111

[39] Hirsch, M. G. et al. Stochastic modeling of single-cell gene expression adaptation reveals non-genomic contribution to evolution of tumor subclones. Cell Syst. 16, 101156 (2025). DOI: 10.1016/j.cels.2024.11.013

[40] Hilfinger, A. & Paulsson, J. Separating intrinsic from extrinsic fluctuations in dynamic biological systems. Proc. Natl Acad. Sci. USA 108, 12167–12172 (2011). DOI: 10.1073/pnas.1018832108

[41] Piras, V., Tomita, M. & Selvarajoo, K. Is central dogma a global property of cellular information flow*?* Front. Physiol. 3, 439 (2012). DOI: 10.3389/fphys.2012.00439

[42] Peccoud, J. & Ycart, B. Markovian modeling of gene-product synthesis. Theor. Popul. Biol. 48, 222–234 (1995). DOI: 10.1006/tpbi.1995.1027

[43] Thattai, M. & van Oudenaarden, A. Intrinsic noise in gene regulatory networks. Proc. Natl Acad. Sci. USA 98, 8614–8619 (2001). DOI: 10.1073/pnas.151588598

[44] Lin, C. Y. et al. Transcriptional amplification in tumor cells with elevated c-Myc. Cell 151, 56–67 (2012). DOI: 10.1016/j.cell.2012.08.026

[45] Friedel, C. C., Dölken, L., Ruzsics, Z., Koszinowski, U. H. & Zimmer, R. Conserved principles of mammalian transcriptional regulation revealed by RNA half-life. Nucleic Acids Res. 37, e115 (2009). DOI: 10.1093/nar/gkp542

[46] Kosik, K. S. Life at low copy number: how dendrites manage with so few mRNAs. Neuron 92, 1168–1180 (2016). DOI: 10.1016/j.neuron.2016.11.002

[47] Charan K, Kar S. Subtle alteration in transcriptional memory governs the lineage-level cell cycle duration heterogeneities of mammalian cells. iScience. 2025 Jun 21;28(7):112981. doi: 10.1016/j.isci.2025.112981.

[48] Bai J, Wu K, Cao MH, Yang Y, Pan Y, Liu H, He Y, Itahana Y, Huang L, Zheng JN, Pan ZQ. SCF^FBXO22^ targets HDM2 for degradation and modulates breast cancer cell invasion and metastasis. Proc Natl Acad Sci U S A. 2019 Jun 11;116(24):11754–11763. doi: 10.1073/pnas.1820990116.

[49] Liu, X. M., Xie, H. Z., Liu, L. G., & Li, Z. B. (2009). Effect of multiplicative and additive noise on genetic transcriptional regulatory mechanism. Physica A: Statistical Mechanics and its Applications, 388(4), 392–398.

[50] Sassi AS, Garcia-Alcala M, Aldana M, Tu Y. Protein concentration fluctuations in the high expression regime: Taylor’s law and its mechanistic origin. Phys Rev X. 2022 Jan-Mar;12(1):011051. DOI: 10.1103/physrevx.12.011051.

[51] Ghosh, S., Banerjee, S. & Bose, I. Emergent bistability: Effects of additive and multiplicative noise. Eur. Phys. J. E 35, 11 (2012). DOI: 10.1140/epje/i2012-12011-4

[52] Zhang Z, Zabaikina I, Nieto C, Vahdat Z, Bokes P, et al. (2025) Stochastic gene expression in proliferating cells: Differing noise intensity in single-cell and population perspectives. PLOS Computational Biology 21(6): e1013014. 10.1371/journal.pcbi.1013014

[53] Ritchie, M. E. et al. limma powers differential expression analyses for RNA-sequencing and microarray studies. Nucleic Acids Res. 43, e47 (2015). DOI: 10.1093/nar/gkv007

[54] Yu, G., Wang, L., Han, Y. & He, Q. clusterProfiler: an R package for comparing biological themes among gene clusters. OMICS 16, 284–287 (2012). DOI: 10.1089/omi.2011.0118

[55] Hayashi, K., Piras, V., Tabata, S. et al. A systems biology approach to suppress TNF-induced proinflammatory gene expressions. Cell Commun Signal 11, 84 (2013). DOI: 10.1186/1478-811X-11-84

[56] Bui, T.T., Selvarajoo, K. Attractor Concepts to Evaluate the Transcriptome-wide Dynamics Guiding Anaerobic to Aerobic State Transition in *Escherichia coli*. Sci Rep 10, 5878 (2020). DOI: 10.1038/s41598-020-62804-3

[57] Gary Wilk, Rosemary Braun, Integrative analysis reveals disrupted pathways regulated by microRNAs in cancer, Nucleic Acids Research, Volume 46, Issue 3, 16 February 2018, Pages 1089–1101, DOI: 10.1093/nar/gkx1250

[58] Battich, N., Stoeger, T. & Pelkmans, L. Control of transcript variability in single mammalian cells. Cell 163, 1596–1610 (2015). DOI: 10.1016/j.cell.2015.11.018

[59] Hayes, J. D., Flanagan, J. U. & Jowsey, I. R. Glutathione transferases. Annu. Rev. Pharmacol. Toxicol. 45, 51–88 (2005). DOI: 10.1146/annurev.pharmtox.45.120403.095857

[60] Edenberg, H. J. & McClintick, J. N. Alcohol dehydrogenases, aldehyde dehydrogenases, and alcohol use disorders: a critical review. Alcohol. Clin. Exp. Res. 42, 2281–2297 (2018). DOI: 10.1111/acer.13904

[61] Horani, A. & Ferkol, T. W. Advances in the genetics of primary ciliary dyskinesia: clinical implications. Chest 154, 645–652 (2018). DOI: 10.1016/j.chest.2018.05.007

[62] Knowles, M. R., Zariwala, M. & Leigh, M. Primary ciliary dyskinesia. Clin. Chest Med. 37, 449–461 (2016). DOI: 10.1016/j.ccm.2016.04.008

[63] Serna, M., Giles, J., Morgan, B. et al. Structural basis of complement membrane attack complex formation. Nat. Commun. 7, 10587 (2016). DOI: 10.1038/ncomms10587

[64] Niwa, S., Nakajima, K., Miki, H., Minato, Y., Wang, D. & Hirokawa, N. KIF19A is a microtubule-depolymerizing kinesin for ciliary length control. Dev. Cell 23, 1167–1175 (2012). DOI: 10.1016/j.devcel.2012.10.016

[65] Xie, N., Zhang, L., Gao, W. et al. NAD+ metabolism: pathophysiologic mechanisms and therapeutic potential. Signal Transduct. Target. Ther. 5, 227 (2020). DOI: 10.1038/s41392-020-00311-7

[66] Barabási, A.-L. & Albert, R. Emergence of scaling in random networks. Science 286, 509–512 (1999). DOI: 10.1126/science.286.5439.509

[67] Yepuri, G., Ramirez, L.M., Theophall, G.G. et al. DIAPH1-MFN2 interaction regulates mitochondria-SR/ER contact and modulates ischemic/hypoxic stress. Nat Commun 14, 6900 (2023). DOI: 10.1038/s41467-023-42521-x

[68] Kwon, J., Kang, J., Jo, A. et al. Single-cell mapping of combinatorial target antigens for CAR switches using logic gates. Nat Biotechnol 41, 1593–1605 (2023). DOI: 10.1038/s41587-023-01686-y

[69] Melouane, A., Ghanemi, A., Aubé, S., Yoshioka, M. & St-Amand, J. Differential gene expression analysis in ageing muscle and drug discovery perspectives. Ageing Res. Rev. 41, 53–63 (2018). DOI: 10.1016/j.arr.2017.10.006

[70] Ogino, S., Fuchs, C. S. & Giovannucci, E. How many molecular subtypes? Implications of the unique tumor principle in personalized medicine. Expert Rev. Mol. Diagn. 12, 621–628 (2012). DOI: 10.1586/erm.12.46

[71] Mroz, E. A. & Rocco, J. W. The challenges of tumor genetic diversity. Cancer 123, 917–927 (2017). DOI: 10.1002/cncr.30430

[72] Kanwal, R. & Gupta, S. Epigenetic modifications in cancer. Clin. Genet. 81, 303–311 (2012). DOI: 10.1111/j.1399-0004.2011.01809.x

[73] Whiteside, T. L. The tumor microenvironment and its role in promoting tumor growth. Oncogene 27, 5904–5912 (2008). DOI: 10.1038/onc.2008.271

[74] Delignette-Muller, M. L. & Dutang, C. fitdistrplus: an R package for fitting distributions. J. Stat. Softw. 64, 1–34 (2015). DOI: 10.18637/jss.v064.i04

[75] Becker, R. A., Chambers, J. M. & Wilks, A. R. The New S Language (Wadsworth & Brooks/Cole, 1988).

[76] Kendall, M. G. A new measure of rank correlation. Biometrika 30, 81–93 (1938). DOI: 10.1093/biomet/30.1-2.81

[77] Kendall, M. G. The treatment of ties in rank problems. Biometrika 33, 239–251 (1945). DOI: 10.1093/biomet/33.3.239

[78] Hausser, J. & Strimmer, K. Entropy inference and the James-Stein estimator, with application to nonlinear gene association networks. J. Mach. Learn. Res. 10, 1469–1484 (2009). [arXiv:0811.3579]

[79] Otasek, D., Morris, J. H., Bouças, J., Pico, A. R. & Demchak, B. Cytoscape automation: empowering workflow-based network analysis. Genome Biol. 20, 185 (2019). DOI: 10.1186/s13059-019-1758-4

[80] Doncheva, N. T., Morris, J. H., Gorodkin, J. & Jensen, L. J. Cytoscape StringApp: network analysis and visualization of proteomics data. J. Proteome Res. 18, 623–632 (2019). DOI: 10.1021/acs.jproteome.8b00702

[81] Doncheva, N. T., Assenov, Y., Domingues, F. S. & Albrecht, M. Topological analysis and interactive visualization of biological networks and protein structures. Nat. Protoc. 7, 670–685 (2012). DOI: 10.1038/nprot.2012.004

