## Supplementary Materials for "Synthetic Data to Explore Transcriptional Regulation of Differentially Expressed Genes in Ovarian Cancer"

### Supplementary Information
### (Figures and Tables)

**Synthetic Data to Explore Transcriptional Regulation of Differentially Expressed Genes in Ovarian Cancer**

Zhang Shuo[1,2] and Kumar Selvarajoo[1,2,3,4*]

[1]School of Biological Sciences, Nanyang Technological University (NTU), 60 Nanyang Drive, SBS-01s-45, Singapore, 637551, Republic of Singapore

[2]Bioinformatics Institute (BII), Agency for Science, Technology and Research (A*STAR), 30 Biopolis Street, Matrix #07-01, Singapore, 138761, Republic of Singapore

[3]Synthetic Biology Translational Research Program, Yong Loo Lin School of Medicine, National University of Singapore (NUS), 10 Medical drive, Singapore, 117597, Republic of Singapore

[4]Synthetic Biology for Clinical and Technological Innovation (SynCTI), National University of Singapore (NUS), 28 Medical Drive, Centre for Life Sciences #02-07, Singapore, 117456, Republic of Singapore

#### Supplementary Figures

**Figure S1. The distributions (histograms) and log-normal fitting for all samples.**

Upper table: The distributions of gene expression for 5 FTEs (blue) and 6 HSGOCs (red). To better visualize the distributions, the gene expression and frequency were taken on a log scale.

Lower table: Besides, the log-normal fitting of gene expression for all samples (light blue) were displayed and the gene expression were taken on a log scale.

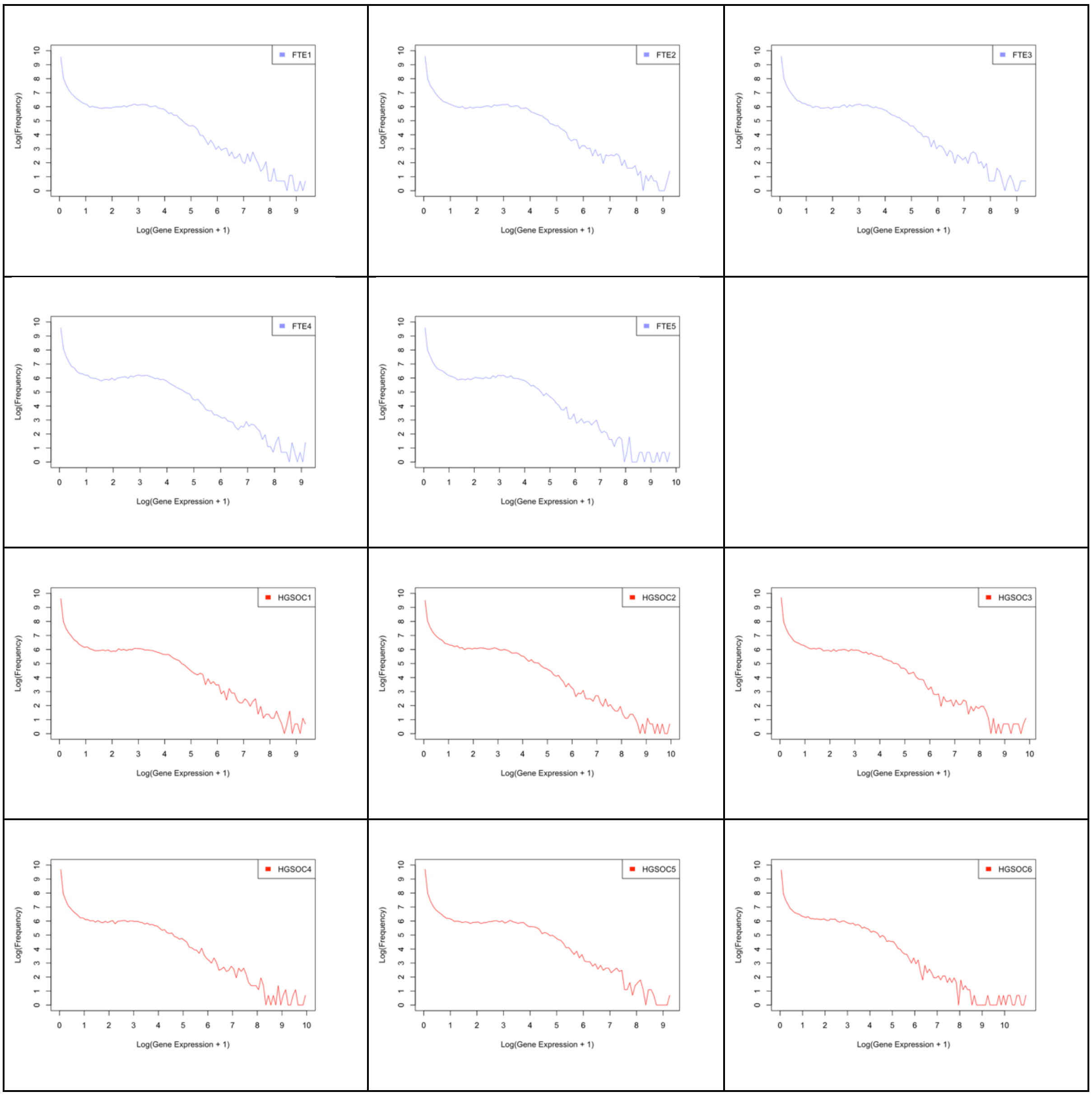

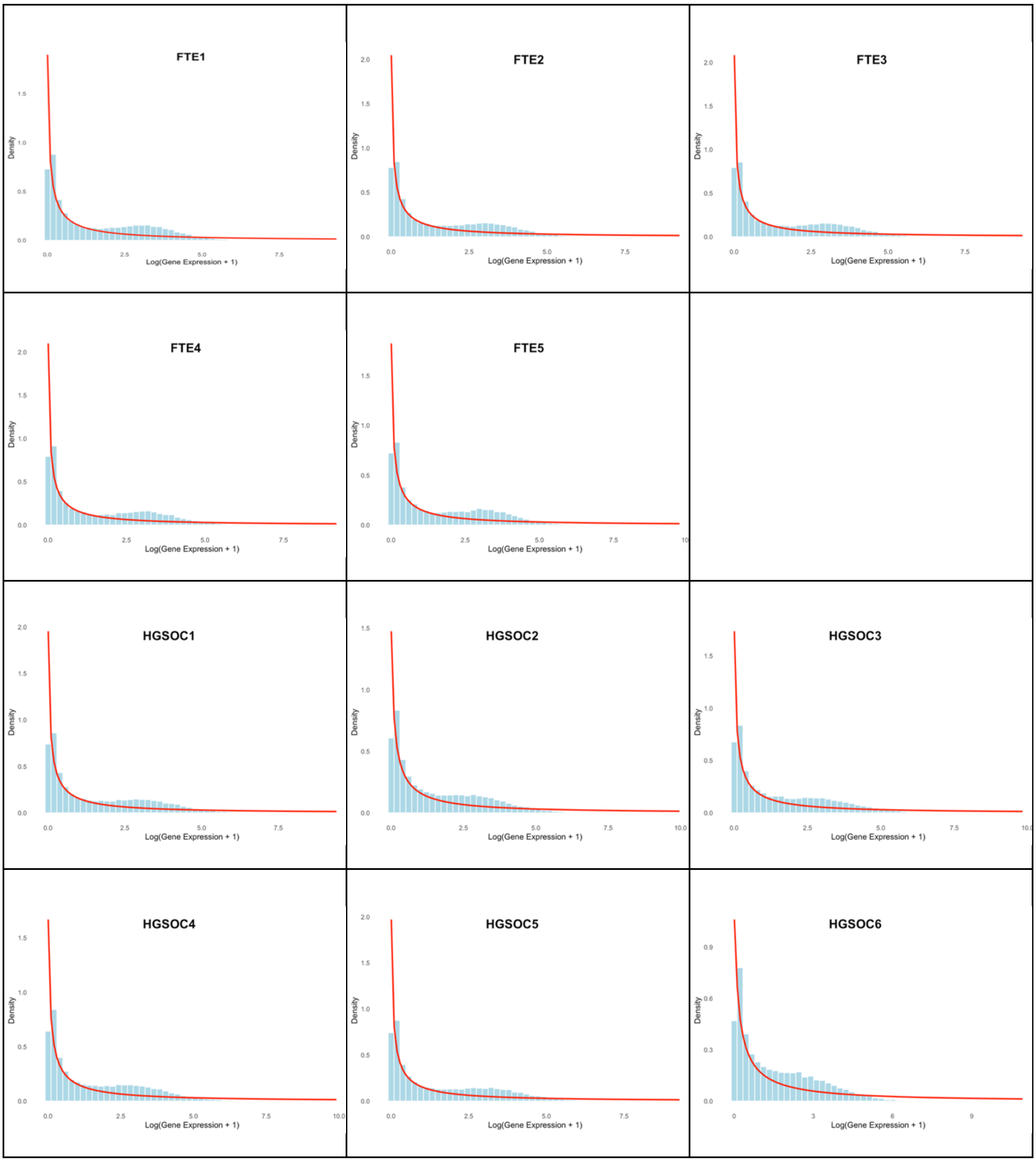
FTE1
FTE2
FTE3
FTE4
FTE5
HGSOC1
HGSOC2
HGSOC3
HGSOC4
HGSOC5
HGSOC6
Density
Log(Gene Expression + 1)

**Figure S2. The transcriptome-wide scatter plots of all samples**

The transcriptome-wide scatter plots for all FTEs (blue) and HSGOCs (red). To better visualize the scatter plots, the scatter plots were taken on a log scale.

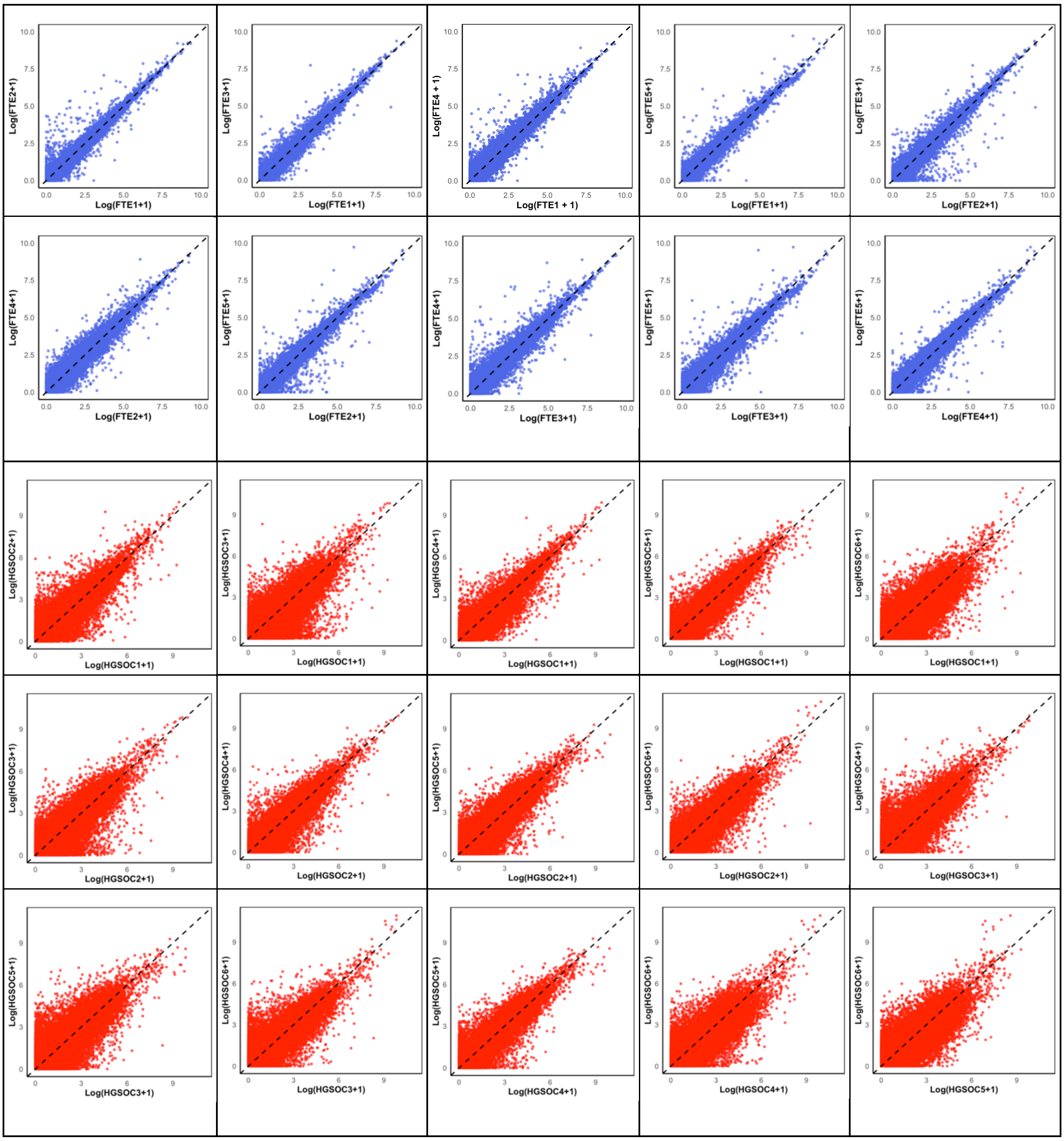

**Figure S3. The comparison of distributions of real distributions and stochastic noises with different groups for Gaussian noises.**

The Gaussian noises $\varphi(z)$ are generated from our method by adjusting the coefficients with $\mu$ and $\sigma$. For instance, we generated $\varphi(z)$ by four groups of $\mu$ and $\sigma$ and plotted their distributions respectively (orange, green, pink and light blue) and compared them to the real distribution (blue). The distributions of stochastic noise (Sn) with Gaussian noises (Gn) generated from the standard normal distribution do not align well with the real distributions.

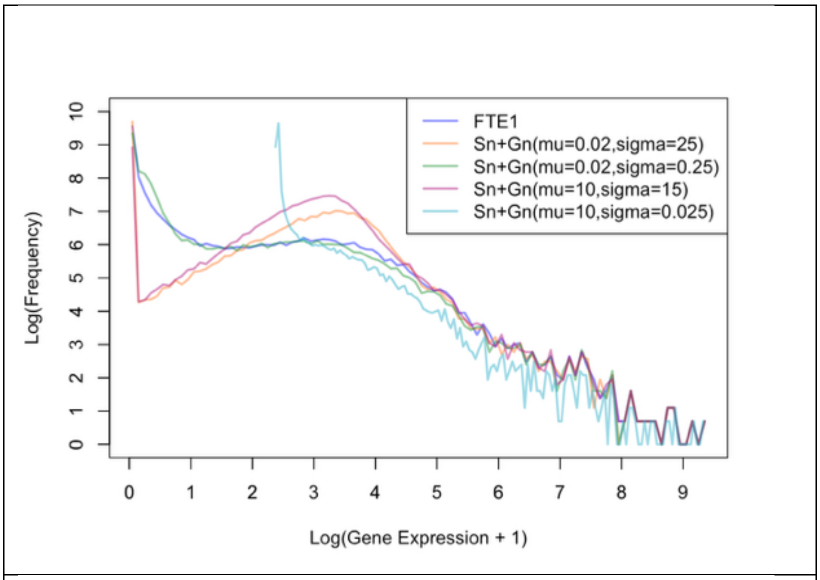

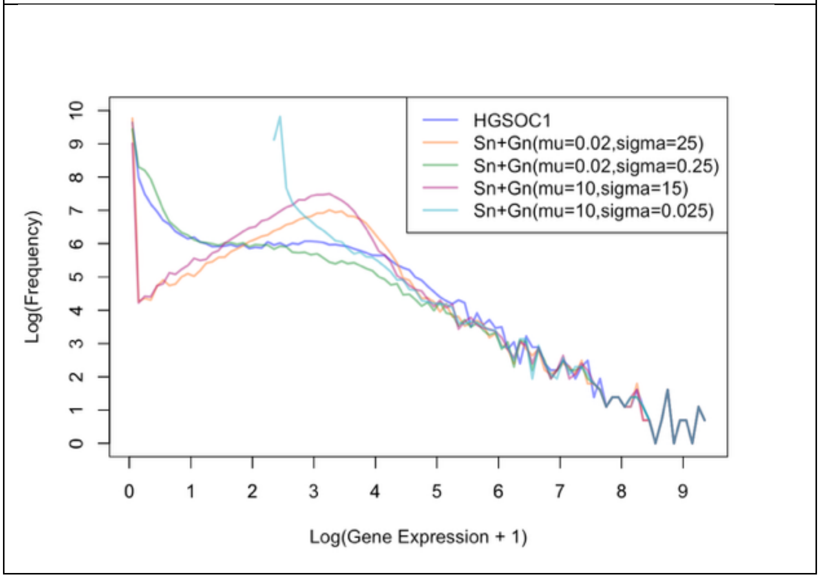

**Figure S4. The comparisons of distribution plots, transcriptome-wide scatter plots (with Gaussian noises) between all experiments and synthetics.**

The comparisons of distributions of gene expression and transcriptome-wide scatter plots between synthetic data (Orange for FTEs & Green for HGSOCs) and experimental data (Blue for FTEs & Red for HGSOCs).

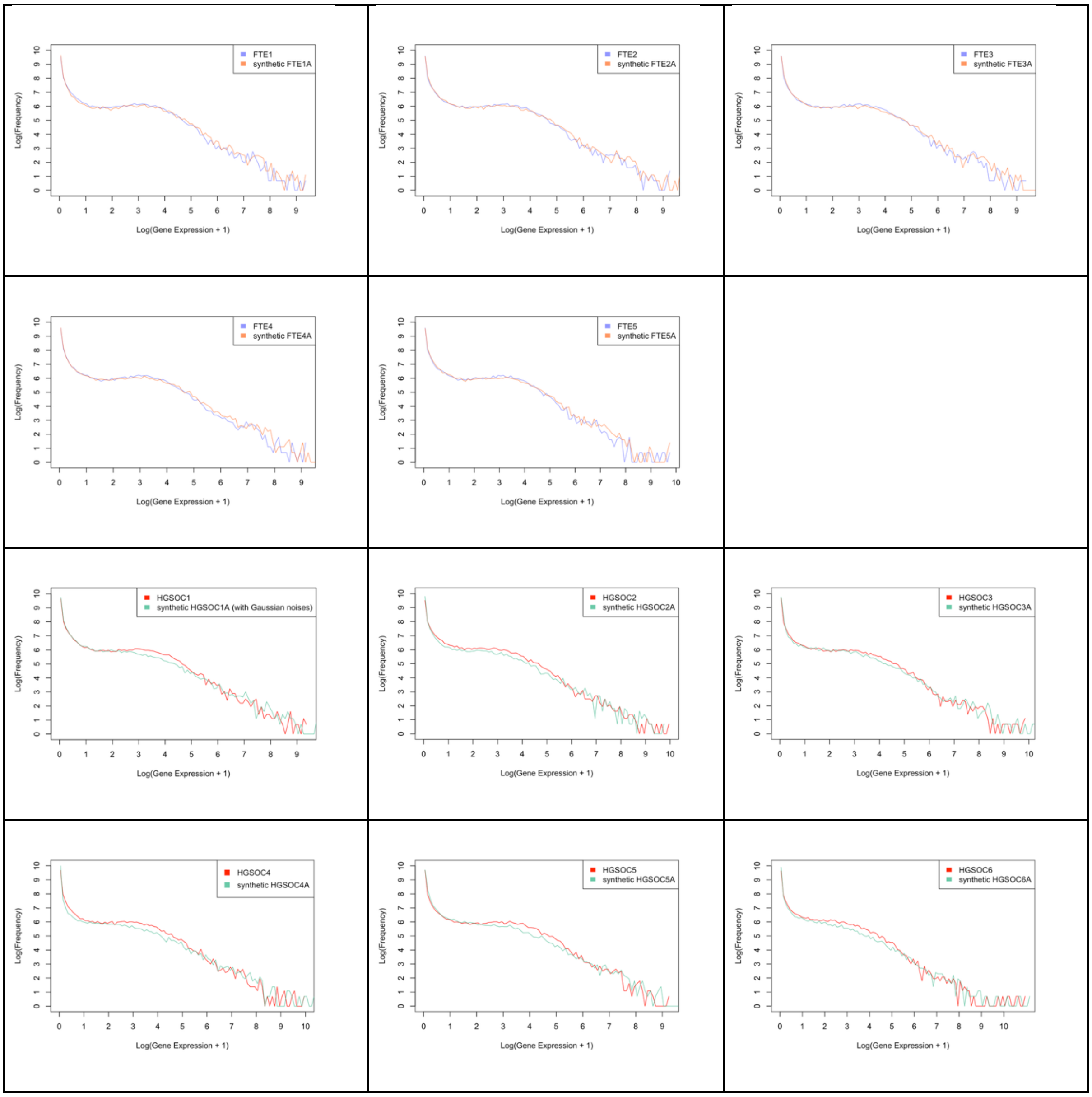

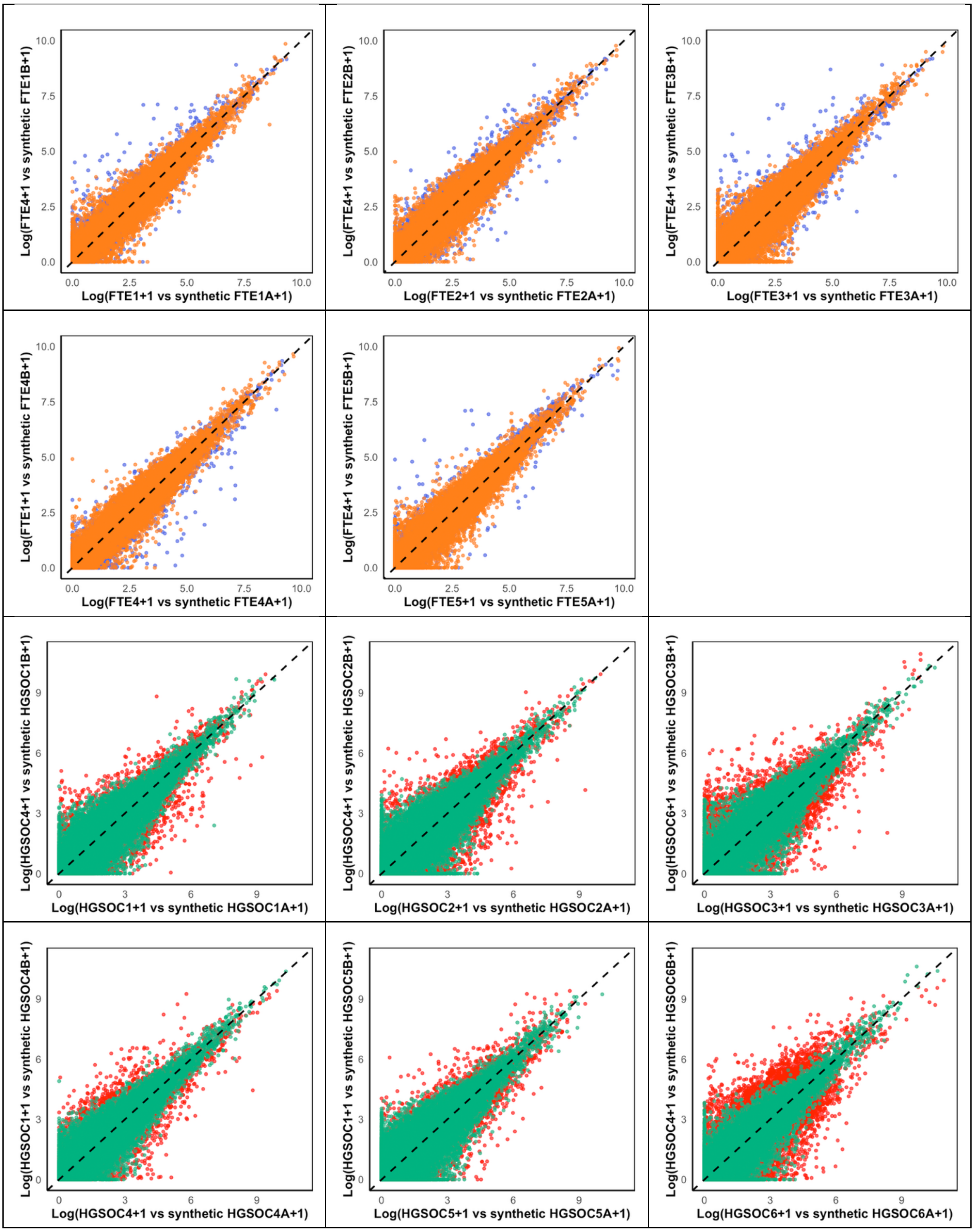
Log(FTE4+1 vs synthetic FTE1B+1)
Log(FTE1+1 vs synthetic FTE1A+1)
Log(FTE4+1 vs synthetic FTE2B+1)
Log(FTE2+1 vs synthetic FTE2A+1)
Log(FTE4+1 vs synthetic FTE3B+1)
Log(FTE3+1 vs synthetic FTE3A+1)
Log(FTE1+1 vs synthetic FTE4B+1)
Log(FTE4+1 vs synthetic FTE4A+1)
Log(FTE4+1 vs synthetic FTE5B+1)
Log(FTE5+1 vs synthetic FTE5A+1)
0.0 2.5 5.0 7.5 10.0
Log(HGSOC4+1 vs synthetic HGSOC1B+1)
Log(HGSOC1+1 vs synthetic HGSOC1A+1)
Log(HGSOC4+1 vs synthetic HGSOC2B+1)
Log(HGSOC2+1 vs synthetic HGSOC2A+1)
Log(HGSOC6+1 vs synthetic HGSOC3B+1)
Log(HGSOC3+1 vs synthetic HGSOC3A+1)
Log(HGSOC1+1 vs synthetic HGSOC4B+1)
Log(HGSOC4+1 vs synthetic HGSOC4A+1)
Log(HGSOC1+1 vs synthetic HGSOC5B+1)
Log(HGSOC5+1 vs synthetic HGSOC5A+1)
Log(HGSOC4+1 vs synthetic HGSOC6B+1)
Log(HGSOC6+1 vs synthetic HGSOC6A+1)
0 3 6 9

**Figure S5. PC2 loading genes removal.**

Removing top 100 high PC2 loading genes to investigate the variability caused by the high PC2 loading genes. After removing the top 2400 PC2 loading genes, our experimental HGSOCs showed 3 distinct clusters, where we require 1 TR model for each of the cluster to better represent the inherent variability of the HGSOCs. (Only steps of 500 genes removed is shown below).

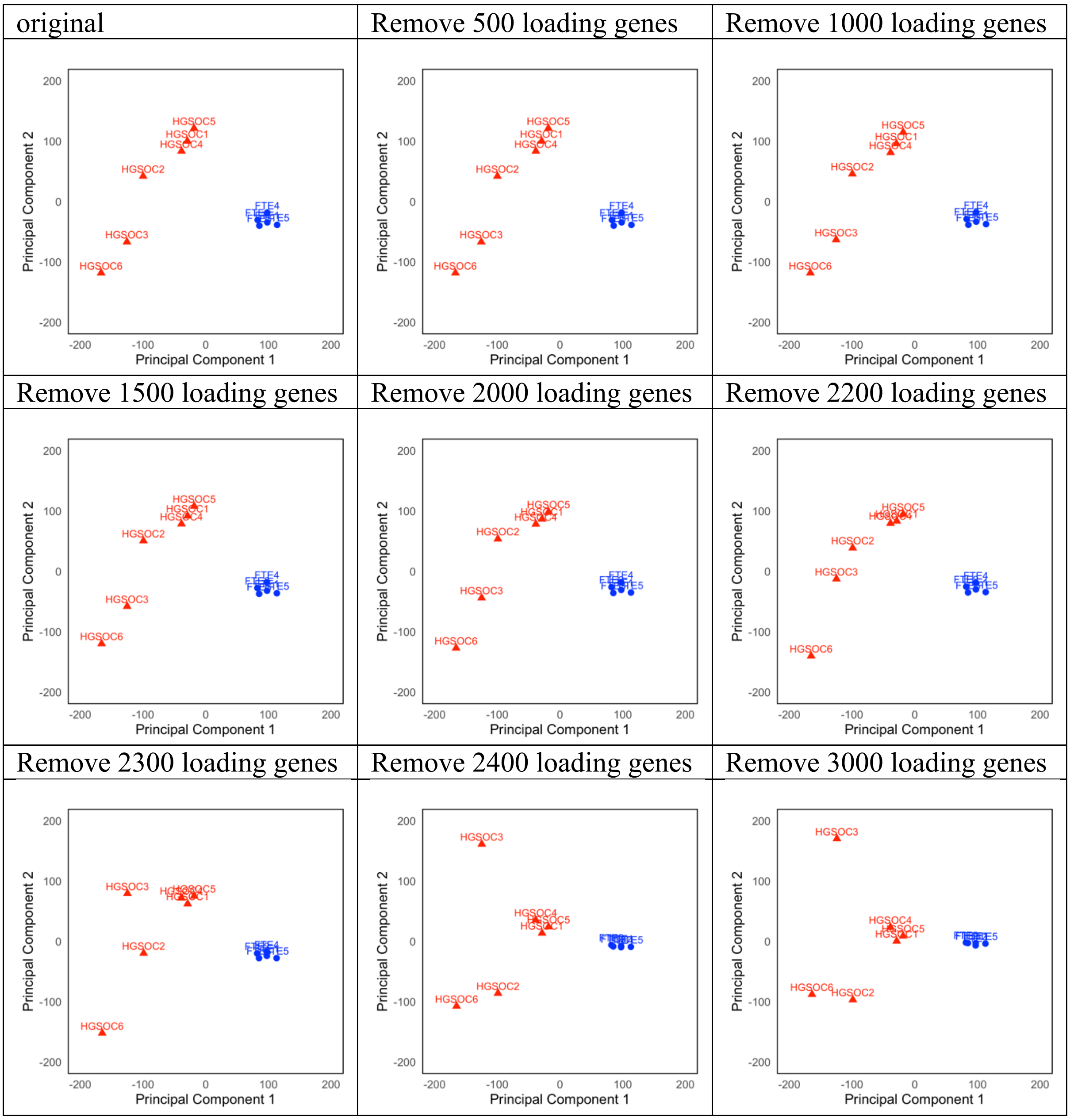

**Figure S6. Stochastic TR model application in the Cardiomyocytes dataset and its scatter plots.**

The stochastic TR model was applied to a normalized Cardiomyocytes dataset (using CPM normalization) and successfully generated comparable synthetic data for the Diaphanous (DIAPH) knockdown and scramble (Scr) control conditions. The corresponding histograms and scatter plots are shown below.

| | Real | Overlap | Distribution |
|---|---|---|---|
| DIAPH | 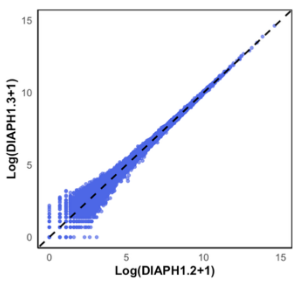  | 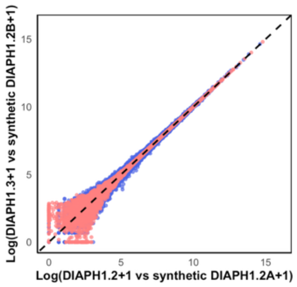  | 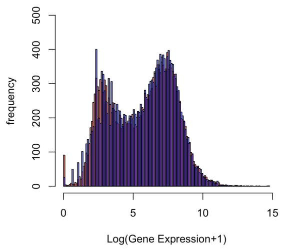  |
| Scr | 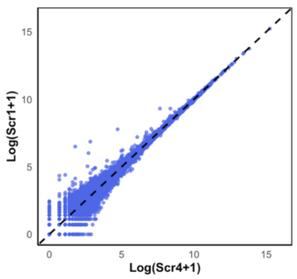  | 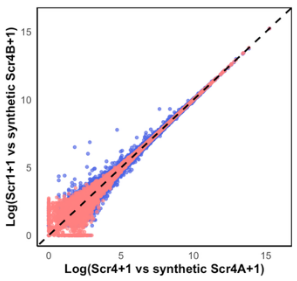  | 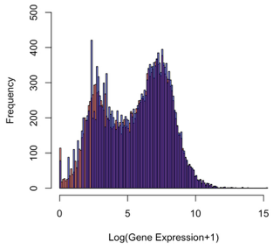  |

**Figure S7. Stochastic TR model application in the top 2000 genes of single cell RNA-Sequencing dataset of ovarian cancer and scatter plots.**

The stochastic TR model was applied to a normalized single-cell RNA sequencing dataset of ovarian cancer. Additionally, we successfully generated comparable synthetic data for the top 2,000 genes, with the corresponding histograms and scatter plots shown below.

<table>
<tr><th></th><th>synthetics</th><th>Overlap</th><th>Distribution</th></tr>
<tr><td>Single cell RNA-Sequencing</td><td>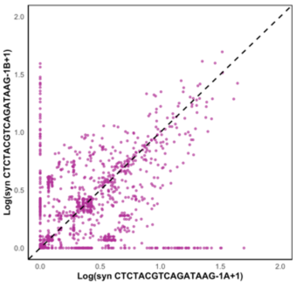
</td><td>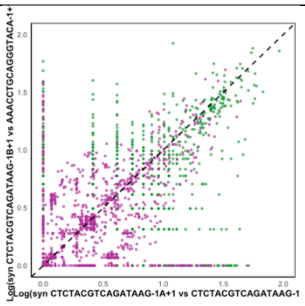
</td><td>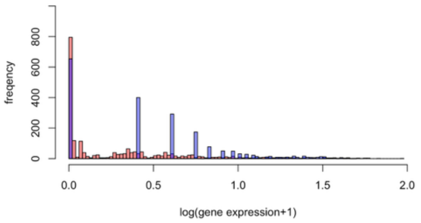
</td></tr>
</table>

**Table S1: Parameter table for the top 100 De genes.**
Parameter values of Kon, Koff, amplification, degradation and transcriptional rate of the top 100 DE genes between FTE and HGSOC replicates from the ovarian cancer cell datasets.

| | Top DE genes | Sample | Kon | Koff | Amplification | Transcription | Degradation |
|---|---|---|---|---|---|---|---|
| 1 | GSTA3 | FTE | 90 | 30 | 5000 | 1.19659810506367 | 0.0994304051718323 |
| | (ENSG00000174156) | HGSOC | 85 | 2 | 4000 | 0.0756515827290036 | 0.310487331623876 |
| 2 | SNTN | FTE | 90 | 30 | 5000 | 22.9905521983924 | 0.716776248094733 |
| | (ENSG00000188817) | HGSOC | 85 | 2 | 4000 | 5.53475245076014 | 1.8788565701323 |
| 3 | ADH1B | FTE | 90 | 30 | 5000 | 0.830903260356058 | 0.261211751675833 |
| | (ENSG00000196616) | HGSOC | 85 | 2 | 4000 | 1.9767881454348 | 0.635017467947457 |
| 4 | DNAI2 | FTE | 90 | 30 | 5000 | 72.383610199025 | 0.59341806657437 |
| | (ENSG00000171595) | HGSOC | 85 | 2 | 4000 | 0.941182687731571 | 2.60289484392773 |
| 5 | CIBAR2 | FTE | 90 | 30 | 5000 | 15.8047732464164 | 0.323663111667644 |
| | (ENSG00000153789) | HGSOC | 85 | 2 | 4000 | 2.23860236395428 | 0.204672384283325 |
| 6 | FAM216B | FTE | 90 | 30 | 5000 | 16.850114098021 | 0.128910482463513 |
| | (ENSG00000179813) | HGSOC | 85 | 2 | 4000 | 4.65810021684203 | 1.58329961688274 |
| 7 | C11orf97 | FTE | 90 | 30 | 5000 | 1.61409256240337 | 0.767697490809119 |
| | (ENSG00000257057) | HGSOC | 85 | 2 | 4000 | 0.00986145436294673 | 0.742090823686628 |
| 8 | ANKRD66 | FTE | 90 | 30 | 5000 | 0.640095255046316 | 0.213829314815428 |
| | (ENSG00000230062) | HGSOC | 85 | 2 | 4000 | 0.230264112662962 | 0.76909452357475 |
| 9 | C6 | FTE | 90 | 30 | 5000 | 1.43165462086541 | 0.160784307675493 |
| | (ENSG00000039537) | HGSOC | 85 | 2 | 4000 | 1.0961353539592 | 0.303396458985068 |
| 10 | KIF19 | FTE | 90 | 30 | 5000 | 12.9729260385517 | 0.275214297959054 |
| | (ENSG00000196169) | HGSOC | 85 | 2 | 4000 | 2.63503096244163 | 0.269775181145016 |
| 11 | LINC01571 | FTE | 90 | 30 | 5000 | 0.070136290212845 | 0.309530935308244 |
| | (ENSG00000260057) | HGSOC | 85 | 2 | 4000 | 2.02738183586825 | 0.406790060527839 |
| 12 | SPAG6 | FTE | 90 | 30 | 5000 | 1.55784295775666 | 0.259434335932861 |
| | (ENSG00000077327) | HGSOC | 85 | 2 | 4000 | 3.63322699962528 | 0.170997184896805 |
| 13 | NDNF | FTE | 90 | 30 | 5000 | 0.0319595341170657 | 0.212536165801484 |
| | (ENSG00000173376) | HGSOC | 85 | 2 | 4000 | 0.123918874931808 | 6.72197705884138 |
| 14 | HOATZ | FTE | 90 | 30 | 5000 | 699.805913390922 | 0.797806766467258 |
| | (ENSG00000183644) | HGSOC | 85 | 2 | 4000 | 0.393060041363092 | 0.303975149683142 |
| 15 | ANKUB1 | FTE | 90 | 30 | 5000 | 1.03014805400177 | 0.627585015631047 |
| | (ENSG00000206199) | HGSOC | 85 | 2 | 4000 | 3.27523241669494 | 0.171370946572117 |
| 16 | CFAP65 | FTE | 90 | 30 | 5000 | 1285.86358525157 | 0.485880871194548 |
| | (ENSG00000181378) | HGSOC | 85 | 2 | 4000 | 0.00935761460794122 | 1.29407062561545 |
| 17 | DTHD1 | FTE | 90 | 30 | 5000 | 9.90414555731559 | 1.88073855832239 |
| | (ENSG00000197057) | HGSOC | 85 | 2 | 4000 | 0.101788462842573 | 0.187579039482963 |
| 18 | LRRC18 | FTE | 90 | 30 | 5000 | 3.26498133629436 | 0.286711014185666 |
| | (ENSG00000165383) | HGSOC | 85 | 2 | 4000 | 0.0653522801995012 | 1.09553562567985 |
| 19 | CACNG6 | FTE | 90 | 30 | 5000 | 0.841232352950713 | 0.481874131615995 |
| | (ENSG00000130433) | HGSOC | 85 | 2 | 4000 | 0.367275860600421 | 0.290280556220658 |
| 20 | C1orf87 | FTE | 90 | 30 | 5000 | 0.0544207989226869 | 0.374315181472066 |
| | (ENSG00000162598) | HGSOC | 85 | 2 | 4000 | 0.333369025625043 | 2.2267711742542 |
| 21 | ERICH3 | FTE | 90 | 30 | 5000 | 1.57213722156543 | 0.258853655083087 |
| | (ENSG00000178965) | HGSOC | 85 | 2 | 4000 | 0.0909682323256049 | 0.601386637988795 |
| 22 | SRD5A2 | FTE | 90 | 30 | 5000 | 2.74893919602196 | 0.12051620920584 |
| | (ENSG00000049319) | HGSOC | 85 | 2 | 4000 | 0.0909724517146826 | 1.57821458188058 |
| 23 | SERPINA6 | FTE | 90 | 30 | 5000 | 4.46123821869457 | 1.64053233808623 |
| | (ENSG00000170099) | HGSOC | 85 | 2 | 4000 | 0.00244801119666795 | 0.640103216575025 |
| 24 | CFAP52 | FTE | 90 | 30 | 5000 | 4.97989047345936 | 0.435812112211033 |
| | (ENSG00000166596) | HGSOC | 85 | 2 | 4000 | 0.707010904960249 | 2.77697353041859 |
| 25 | ABCA8 | FTE | 90 | 30 | 5000 | 1.71670010792313 | 0.360609247768433 |
| | (ENSG00000141338) | HGSOC | 85 | 2 | 4000 | 0.738460862299941 | 1.36528807094501 |
| 26 | PPIAP45 | FTE | 90 | 30 | 5000 | 5.10081134989435 | 0.0957288140819307 |
| | (ENSG00000258116) | HGSOC | 85 | 2 | 4000 | 15.2247855350423 | 0.794391626730196 |
| 27 | SMOC1 | FTE | 90 | 30 | 5000 | 3.88820760545465 | 0.10153786139427 |
| | (ENSG00000198732) | HGSOC | 85 | 2 | 4000 | 0.274718551326915 | 1.75816626900616 |
| 28 | DPT | FTE | 90 | 30 | 5000 | 0.186825300800875 | 0.141959556585384 |
| | (ENSG00000143196) | HGSOC | 85 | 2 | 4000 | 3.29971785169947 | 0.334660729027247 |
| 29 | UMODL1-AS1 | FTE | 90 | 30 | 5000 | 5.64799493275684 | 3.69735138263532 |
| | ENSG00000184385 | HGSOC | 85 | 2 | 4000 | 12.8509493529913 | 2.20633161305879 |
| 30 | DNAH9 | FTE | 90 | 30 | 5000 | 1493.21231005452 | 0.356832456292576 |
| | ENSG00000007174 | HGSOC | 85 | 2 | 4000 | 3.2399583776162 | 0.159660167065688 |

| | | | | | | | |
|---|---|---|---|---|---|---|---|
| 31 | RP11-268F1.3 | FTE | 90 | 30 | 5000 | 1.5373909751379 | 0.473218858141577 |
| | (ENSG00000236164) | HGSOC | 85 | 2 | 4000 | 0.100052715712733 | 1.10516642304108 |
| 32 | SLC23A1 | FTE | 90 | 30 | 5000 | 8.44193961955931 | 0.308262719944821 |
| | (ENSG00000170482) | HGSOC | 85 | 2 | 4000 | 2.01615217771499 | 0.173827604470175 |
| 33 | CRISP3 | FTE | 90 | 30 | 5000 | 0.362893342419722 | 0.109034435838987 |
| | (ENSG00000096006) | HGSOC | 85 | 2 | 4000 | 29.7597792781739 | 1.69258826174056 |
| 34 | PRR29 | FTE | 90 | 30 | 5000 | 0.273753644785778 | 0.159283424655468 |
| | (ENSG00000224383) | HGSOC | 85 | 2 | 4000 | 3.96359245780444 | 1.2788275578171 |
| 35 | PPP1R42 | FTE | 90 | 30 | 5000 | 3.30073904658607 | 0.876613417180499 |
| | (ENSG00000178125) | HGSOC | 85 | 2 | 4000 | 0.118280097641478 | 0.452486392550859 |
| 36 | CCL21 | FTE | 90 | 30 | 5000 | 1.09045302249251 | 0.065972305069186 |
| | (ENSG00000137077) | HGSOC | 85 | 2 | 4000 | 2.72924571021078 | 1.61044415622069 |
| 37 | SFRP1 | FTE | 90 | 30 | 5000 | 0.0827702798852884 | 0.195845293743537 |
| | (ENSG00000104332) | HGSOC | 85 | 2 | 4000 | 3.05728276285041 | 0.216759326102522 |
| 38 | WDR38 | FTE | 90 | 30 | 5000 | 1.24605768149593 | 0.284481299396642 |
| | (ENSG00000136918) | HGSOC | 85 | 2 | 4000 | 9.80613378420285 | 0.603910624957529 |
| 39 | APOBEC4 | FTE | 90 | 30 | 5000 | 0.0462998120521006 | 0.173845829749528 |
| | (ENSG00000173627) | HGSOC | 85 | 2 | 4000 | 37.2708365099977 | 0.198315036894897 |
| 40 | MYH11 | FTE | 90 | 30 | 5000 | 11.1585906849658 | 0.25302399701695 |
| | (ENSG00000133392) | HGSOC | 85 | 2 | 4000 | 43.0528206069599 | 0.685970144350629 |
| 41 | CABCOCO1 | FTE | 90 | 30 | 5000 | 0.378234192156183 | 0.35156118392609 |
| | (ENSG00000183346) | HGSOC | 85 | 2 | 4000 | 1.19914847227269 | 0.251749294480616 |
| 42 | ODAD2 | FTE | 90 | 30 | 5000 | 0.232281643937356 | 0.451309807300127 |
| | (ENSG00000169126) | HGSOC | 85 | 2 | 4000 | 0.344621032966774 | 2.53073487740231 |
| 43 | TTC29 | FTE | 90 | 30 | 5000 | 0.695234785219641 | 0.329976524845398 |
| | (ENSG00000137473) | HGSOC | 85 | 2 | 4000 | 0.0830793238305798 | 0.612299211298757 |
| 44 | ROPN1L | FTE | 90 | 30 | 5000 | 0.369683405596189 | 0.562469062765733 |
| | (ENSG00000145491) | HGSOC | 85 | 2 | 4000 | 15.6282027562592 | 0.503082488003682 |
| 45 | AK7 | FTE | 90 | 30 | 5000 | 6.24239386500743 | 0.247923613551723 |
| | (ENSG00000140057) | HGSOC | 85 | 2 | 4000 | 4.01408838114714 | 0.75571905229612 |
| 46 | NHLRC4 | FTE | 90 | 30 | 5000 | 0.259472499567423 | 0.315255309889393 |
| | (ENSG00000257108) | HGSOC | 85 | 2 | 4000 | 0.0637037667413135 | 0.111765695649498 |
| 47 | ESPL1 | FTE | 90 | 30 | 5000 | 0.108912697754314 | 0.174488193322483 |
| | (ENSG00000135476) | HGSOC | 85 | 2 | 4000 | 0.456849214369049 | 0.716067026693878 |
| 48 | LRRC74B | FTE | 90 | 30 | 5000 | 8.72310140977138 | 1.28628713052053 |
| | (ENSG00000187905) | HGSOC | 85 | 2 | 4000 | 0.0944947325491931 | 0.771974559004155 |
| 49 | RP11-445L6.3 | FTE | 90 | 30 | 5000 | 1.15029200885233 | 0.488927250905017 |
| | (ENSG00000234692) | HGSOC | 85 | 2 | 4000 | 55.9360393495593 | 0.302142733052263 |
| 50 | CNGA4 | FTE | 90 | 30 | 5000 | 3.30008810587793 | 0.182878910896538 |
| | (ENSG00000132259) | HGSOC | 85 | 2 | 4000 | 0.367522225206173 | 0.225675616919003 |
| 51 | IQGAP3 | FTE | 90 | 30 | 5000 | 0.69488859367659 | 0.731146240921526 |
| | (ENSG00000183856) | HGSOC | 85 | 2 | 4000 | 0.440421036341462 | 1.5326476387032 |
| 52 | RP11-661D19.3 | FTE | 90 | 30 | 5000 | 0.241490111802907 | 0.523437848583302 |
| | (ENSG00000259508) | HGSOC | 85 | 2 | 4000 | 0.0496818920663274 | 0.237379712225232 |
| 53 | LINC02345 | FTE | 90 | 30 | 5000 | 3.06534370987404 | 0.664515357165718 |
| | (ENSG00000259225) | HGSOC | 85 | 2 | 4000 | 2.97215600142355 | 0.546278743835807 |
| 54 | UBE2C | FTE | 90 | 30 | 5000 | 0.0346087709039482 | 0.361930503498622 |
| | (ENSG00000175063) | HGSOC | 85 | 2 | 4000 | 5.74659106743184 | 1.16089409557377 |
| 55 | AQP10 | FTE | 90 | 30 | 5000 | 5.20099768925572 | 0.499586466819267 |
| | (ENSG00000143595) | HGSOC | 85 | 2 | 4000 | 0.437651102860208 | 3.37360447580452 |
| 56 | CWH43 | FTE | 90 | 30 | 5000 | 0.516269855659333 | 0.177901965873928 |
| | E(NSG00000109182) | HGSOC | 85 | 2 | 4000 | 0.800157948486437 | 3.50490385255424 |
| 57 | C1orf141 | FTE | 90 | 30 | 5000 | 0.689227125524402 | 0.31268704974735 |
| | (ENSG00000203963) | HGSOC | 85 | 2 | 4000 | 0.296278952041929 | 0.838985936233974 |
| 58 | KCNE1 | FTE | 90 | 30 | 5000 | 3.46073115983155 | 0.102136982623454 |
| | (ENSG00000180509) | HGSOC | 85 | 2 | 4000 | 0.758692249953418 | 0.400320837998656 |
| 59 | TTLL6 | FTE | 90 | 30 | 5000 | 1.77275504075043 | 0.141546644065243 |
| | (ENSG00000170703) | HGSOC | 85 | 2 | 4000 | 0.500330947353332 | 10.1583651622252 |
| 60 | HAND2 | FTE | 90 | 30 | 5000 | 0.0635536021678429 | 0.308154330674207 |
| | (ENSG00000164107) | HGSOC | 85 | 2 | 4000 | 0.190239451659324 | 1.39629201093444 |
| 61 | MS4A8 | FTE | 90 | 30 | 5000 | 0.755700630539587 | 0.230700463941332 |
| | (ENSG00000166959) | HGSOC | 85 | 2 | 4000 | 2.27429009193155 | 1.69564905968223 |
| 62 | ALDH1A1 | FTE | 90 | 30 | 5000 | 0.14291968650994 | 0.204384040866396 |
| | (ENSG00000165092) | HGSOC | 85 | 2 | 4000 | 0.574143208520063 | 0.222612784103291 |
| 63 | LRRC71 | FTE | 90 | 30 | 5000 | 6.02423076507822 | 0.343139197518239 |
| | (ENSG00000160838) | HGSOC | 85 | 2 | 4000 | 2.13932508240226 | 0.151307559581105 |
| 64 | CDC20 | FTE | 90 | 30 | 5000 | 3.71611594715294 | 0.947394211429063 |

| | | | | | | | |
|---|---|---|---|---|---|---|---|
| | (ENSG00000117399) | **HGSOC** | **85** | **2** | **4000** | **9.610111475832** | **0.357618821216221** |
| 65 | CCDC81 | **FTE** | **90** | **30** | **5000** | **4.76725600234444** | **0.356184484225086** |
| | (ENSG00000149201) | **HGSOC** | **85** | **2** | **4000** | **5.42339365293247** | **0.667575746479794** |
| 66 | ABCA9 | **FTE** | **90** | **30** | **5000** | **4.73572579826106** | **0.869796236595373** |
| | (ENSG00000154258) | **HGSOC** | **85** | **2** | **4000** | **3.14974023543204** | **0.186968166195196** |
| 67 | SPATA4 | **FTE** | **90** | **30** | **5000** | **1.33613231189155** | **0.756980100264623** |
| | (ENSG00000150628) | **HGSOC** | **85** | **2** | **4000** | **18.0448542845109** | **0.17923715717166** |
| 68 | DNAH3 | **FTE** | **90** | **30** | **5000** | **25.608183347438** | **0.085582076808503** |
| | (ENSG00000158486) | **HGSOC** | **85** | **2** | **4000** | **23.9607413244137** | **0.661595684417082** |
| 69 | DYNLT5 | **FTE** | **90** | **30** | **5000** | **1.10792253502616** | **0.500225488775213** |
| | (ENSG00000152760) | **HGSOC** | **85** | **2** | **4000** | **52.3331716407305** | **1.89684960787969** |
| 70 | CXXC4 | **FTE** | **90** | **30** | **5000** | **0.361817410344986** | **1.20480158050156** |
| | (ENSG00000168772) | **HGSOC** | **85** | **2** | **4000** | **1.89768786152842** | **0.496799618355025** |
| 71 | FMO3 | FTE | **90** | **30** | **5000** | **29.7129740929968** | **0.350933212958044** |
| | (ENSG00000007933) | **HGSOC** | **85** | **2** | **4000** | **19.3397723527248** | **0.109845216734802** |
| 72 | PTPRN2 | **FTE** | **90** | **30** | **5000** | **3.74655439651693** | **0.191567644226941** |
| | (ENSG00000155093) | **HGSOC** | **85** | **2** | **4000** | **0.0579343690064687** | **0.414035002802294** |
| 73 | ADH6 | **FTE** | **90** | **30** | **5000** | **2.56116809094411** | **0.445242913214684** |
| | (ENSG00000172955) | **HGSOC** | **85** | **2** | **4000** | **1.07204410197686** | **1.18654048013967** |
| 74 | CFAP157 | **FTE** | **90** | **30** | **5000** | **0.521578044009014** | **1.17151855992414** |
| | (ENSG00000160401) | **HGSOC** | **85** | **2** | **4000** | **4.34085717983835** | **0.88766637223403** |
| 75 | HAND2-AS1 | **FTE** | **90** | **30** | **5000** | **57.3966011695641** | **0.18566916326837** |
| | (ENSG00000237125) | **HGSOC** | **85** | **2** | **4000** | **0.433997467585924** | **2.56576066437018** |
| 76 | EFHB | **FTE** | **90** | **30** | **5000** | **0.138173902014194** | **1.22606651102027** |
| | (ENSG00000163576) | **HGSOC** | **85** | **2** | **4000** | **0.160818788972003** | **0.341878199929933** |
| 77 | NEK10 | **FTE** | **90** | **30** | **5000** | **0.758231831286521** | **0.398111739955856** |
| | (ENSG00000163491) | **HGSOC** | **85** | **2** | **4000** | **2.72520710879299** | **0.94361612315594** |
| 78 | AGR2 | **FTE** | **90** | **30** | **5000** | **73.0144680675663** | **0.486929821133166** |
| | (ENSG00000106541) | **HGSOC** | **85** | **2** | **4000** | **43.9731132910709** | **0.175607689662428** |
| 79 | RERGL | **FTE** | **90** | **30** | **5000** | **0.0536861795736412** | **0.232916298969554** |
| | (ENSG00000111404) | **HGSOC** | **85** | **2** | **4000** | **0.106550065346276** | **0.937920206457524** |
| 80 | C20orf85 | **FTE** | **90** | **30** | **5000** | **13.4184890874702** | **0.780488080755245** |
| | (ENSG00000124237) | **HGSOC** | **85** | **2** | **4000** | **1.7057727903473** | **1.3645983969957** |
| 81 | OGN | **FTE** | **90** | **30** | **5000** | **226.243504091287** | **0.163135420891514** |
| | (ENSG00000106809) | **HGSOC** | **85** | **2** | **4000** | **0.19225281655505** | **0.976796670140669** |
| 82 | GRCh37 | **FTE** | **90** | **30** | **5000** | **3.49897460932271** | **0.128284684154796** |
| | (ENSG00000253474) | **HGSOC** | **85** | **2** | **4000** | **0.170489208740684** | **0.157759047914462** |
| 83 | RP11-680N20.1 | **FTE** | **90** | **30** | **5000** | **644.773366703343** | **1.16219064949621** |
| | (ENSG00000263450) | **HGSOC** | **85** | **2** | **4000** | **0.0837959980800307** | **0.701145279606183** |
| 84 | DRC3 | **FTE** | **90** | **30** | **5000** | **0.826859912900272** | **0.131510918638896** |
| | (ENSG00000171962) | **HGSOC** | **85** | **2** | **4000** | **0.926195976906563** | **0.221043681546965** |
| 85 | OMG | **FTE** | **90** | **30** | **5000** | **36.7466659027963** | **0.621671711796941** |
| | (ENSG00000126861) | **HGSOC** | **85** | **2** | **4000** | **53.738149240447** | **0.714185308703442** |
| 86 | TCTE1 | **FTE** | **90** | **30** | **5000** | **3.27791096903743** | **0.192722391898551** |
| | (ENSG00000146221) | **HGSOC** | **85** | **2** | **4000** | **0.194154840269309** | **0.533797782319792** |
| 87 | CFAP299 | **FTE** | **90** | **30** | **5000** | **27.3293522685056** | **0.820223653098002** |
| | (ENSG00000197826) | **HGSOC** | **85** | **2** | **4000** | **0.435441809646608** | **0.169389586874093** |
| 88 | NXPH3 | **FTE** | **90** | **30** | **5000** | **0.108421849100147** | **0.465619025227839** |
| | (ENSG00000182575) | **HGSOC** | **85** | **2** | **4000** | **1.52716955702172** | **0.429034933753274** |
| 89 | CFAP61 | **FTE** | **90** | **30** | **5000** | **0.240039156683091** | **0.509612928000471** |
| | (ENSG00000089101) | **HGSOC** | **85** | **2** | **4000** | **4.32528020206593** | **0.598658952781179** |
| 90 | SRGAP3-AS2 | **FTE** | **90** | **30** | **5000** | **24.7062427993972** | **0.084188335192131** |
| | (ENSG00000228723) | **HGSOC** | **85** | **2** | **4000** | **0.313976837737986** | **0.551615505685487** |
| 91 | TROAP | FTE | **90** | **30** | **5000** | **14.9209651318753** | **0.177556853959681** |
| | (ENSG00000135451) | **HGSOC** | **85** | **2** | **4000** | **0.523210621056491** | **1.13811053672256** |
| 92 | FAM166B | **FTE** | **90** | **30** | **5000** | **0.623827654123291** | **0.147250145863211** |
| | (ENSG00000215187) | **HGSOC** | **85** | **2** | **4000** | **1.26770831166126** | **0.193679615202755** |
| 93 | RP11-266I14.3 | **FTE** | **90** | **30** | **5000** | **0.917747416910671** | **0.393118668290409** |
| | (ENSG00000227787) | **HGSOC** | **85** | **2** | **4000** | **1.74906732478701** | **0.719300706138399** |
| 94 | TMEM212 | **FTE** | **90** | **30** | **5000** | **70.2436213968088** | **0.290223724968398** |
| | (ENSG00000186329) | **HGSOC** | **85** | **2** | **4000** | **0.616359561641833** | **0.322103879972841** |
| 95 | PLG | **FTE** | **90** | **30** | **5000** | **11.2646898389484** | **0.390571893661499** |
| | (ENSG00000122194) | **HGSOC** | **85** | **2** | **4000** | **0.556739396575516** | **0.985267804009841** |
| 96 | OPLAH | **FTE** | **90** | **30** | **5000** | **1.76767489407809** | **0.164575743626909** |
| | (ENSG00000178814) | **HGSOC** | **85** | **2** | **4000** | **16.1981027172147** | **0.139523208743754** |
| 97 | SYNE1 | **FTE** | **90** | **30** | **5000** | **0.282073466287373** | **1.33131977865206** |
| | (ENSG00000131018) | **HGSOC** | **85** | **2** | **4000** | **3.32423266657423** | **0.209674036166234** |

| | | | | | | | |
|---|---|---|---|---|---|---|---|
| 98 | Z98881.1 | **FTE** | **90** | **30** | **5000** | **1.6018740073152** | **0.597762663841341** |
| | (ENSG00000196674) | **HGSOC** | **85** | **2** | **4000** | **1.73707037399745** | **0.635527366020099** |
| 99 | CTC-498J12.1 | **FTE** | **90** | **30** | **5000** | **284.295480363855** | **0.932182589056472** |
| | (ENSG00000250237) | **HGSOC** | **85** | **2** | **4000** | **0.228069944916043** | **0.875233533305565** |
| 100 | RP11-664D7.4 | **FTE** | **90** | **30** | **5000** | **0.431846943527987** | **0.027932540927791** |
| | (ENSG00000248801) | **HGSOC** | **85** | **2** | **4000** | **3.53067236622144** | **1.00708072851884** |
